# The influence of new SARS-CoV-2 variant Omicron (B.1.1.529) on vaccine efficacy, its correlation to Delta Variants: a computational approach

**DOI:** 10.1101/2021.12.06.471215

**Authors:** Prashant Ranjan, Neha, Chandra Devi, Kaviyapriya Arulmozhi Devar, Parimal Das

**Author notes:** **Correspondence:** Parimal Das, Centre for Genetic Disorders, Institute of Science, Banaras Hindu University, Varanasi 221005,Uttar Pradesh, India.

## Abstract

The newly discovered COVID variant B.1.1.529 in Botswana has more than 30 mutations in spike and many other in non-spike proteins, far more than any other SARS-CoV-2 variant accepted as a variant of concern by the WHO and officially named Omicron, and has sparked concern among scientists and the general public. Our findings provide insights into structural modification caused by the mutations in the Omicrons receptor-binding domain and look into the effects on interaction with the hosts neutralising antibodies CR3022, B38, CB6, P2B-2F6, and REGN, as well as ACE2R using an *in silico* approach. We have employed secondary structure prediction, structural superimposition, protein disorderness, molecular docking, and MD simulation to investigate host-pathogen interactions, immune evasion, and transmissibility caused by mutations in the RBD region of the spike protein of the Omicron variant and compared it to the Delta variants (AY.1, AY.2, & AY.3) and wild type. Computational analysis revealed that the Omicron variant has a higher binding affinity for the human ACE2 receptor than the wild and Delta (AY.1 and AY.2 strains), but lower than the Delta AY.3 strain. MD simulation and docking analysis suggest that the omicron and Delta AY.3 were found to have relatively unstable and compact RBD structures and hampered interactions with antibodies more than wild and Delta (AY.1 and AY.2), which may lead to relatively more pathogenicity and antibody escape. In addition, we observed lower binding affinity of Omicron for human monoclonal antibodies (CR3022, B38, CB6, and P2B2F6) when compared to wild and Delta (AY.1 & AY.2). However, the binding affinity of Omicron RBD variants for CR3022, B38, and P2B2F6 antibodies is lower as compared to Delta AY.3, which might promote immune evasion and reinfection and needs further experimental investigation.

## 1. Introduction

Viruses naturally have the ability to change their genetic makeup with time, which does not affect it drastically, but these changes may affect host range, disease severity, transmissibility, diagnosis, re-infection, the performance of vaccines and other therapeutics, etc.^1^. The SARS-CoV-2 was first reported in December 2019 in Wuhan, Hubei province, China, where it later became a pandemic^2^. Many variants have emerged since then, causing multiple waves of infection. Among those, Alpha (B.1.1.7 lineage), Beta (B.1.351), Gamma (P.1), and Delta (B.1.617.2) have been categorised as variants of concern (VOCs) by WHO (Campbell et al. 2021). Several vaccines like the Pfizer/BioNTechComirnaty vaccine, SII/Covishield and AstraZeneca/AZD1222 vaccines, Janssen/Ad26.COV 2.S, Moderna COVID-19 vaccine (mRNA 1273), Sinopharm COVID-19 vaccine, and Sinovac-CoronaVac have been deployed for COVID-19 over time. Data from different studies show the effectiveness of vaccines against infection and the severity of the disease caused by the original SARS-CoV-2 strain. However, whether the vaccines are highly effective against other variants is still unknown and requires real-world validation ^3^.

The coronavirus spike (S) protein located throughout the virus surface is important for interaction with the host cell. The S protein (1273aa) is composed of signal peptide (amino acids 1–13), the S1 subunit (14–685aa), and the S2 subunit (686–1273aa). S1 and S2 mainly mediate host angiotensin-converting enzyme-2 (ACE2) receptor recognition and binding, followed by membrane fusion ^4–7^. To be precise, it is the RBD domain (319–541aa residues) located in the S1 subunit that binds to the host cell ACE2 receptor ^6^. After binding with ACE2, a serine protease TMPRSS2 located on the host cell membrane is essentially required for S protein priming and helps viral spread. This TMPRSS2 (transmembrane protease, serine-2) proteolytically cleaves the peptide bond Arg685-Ser686 of the S1/S2 site and separates the two subunits. The S2 subunit, which remains within the viral envelop, ultimately attaches to the host membrane. The process of fusion and infection is further enhanced due to irreversible conformational changes of S protein by the second proteolysis at the S2’ site (Arg815-Ser816). TMPRSS2 also cleaves ACE2 and promotes uptake of SARS-CoV and likely SARS-CoV-2 virions as well. In the secretory pathway of the infected host cell, SARS-CoV-2 S protein is preactivated by furin-mediated proteolysis, which requires a single bond cleavage for the fusion process activation and subsequent entry into the cell ^8,9^.

Mutations in the spike region may affect the way the virus interacts with the host or responds to antibodies. The D614G mutation in spike protein enabled higher ACE2 binding affinity and was correlated with higher transmission and increased viral loads in COVID-19 patients ^7,10^. The Delta variant (B.1.617.2) of SARS-CoV-2 with higher infectivity was the leading factor in the third wave of the COVID-19 pandemic ^11^. Multiple sub-lineages of Delta variants were observed to be circulated among populations. From data submitted in Nextstrain database, the PANGO lineage AY.1 contains T19R, T95I, G142D, E156-, F157-, R158G, W258L, K417N, L452R, T478K, D614G, P681R, D950N and PANGO AY.2 contains T19R, G142D, E156-, F157-, R158G, A222V, K417N, L452R, T478K, D614G, P681R, D950N and PANGO lineage AY.3 contains T19R, E156-, F157-, R158G, L452R, T478K, D614G, P681R, D950N mutations in S protein. The K417N, L452R, and T487K are common mutations in the AY.1 and AY.2 spike RBD, while the K417N mutation is absent in the AY.3 sub-lineage. D614G, P681R, and D950N are other key S protein substitutions in the fusion region present in all Delta sub-lineages. The characteristic Delta variant mutation Del157-158 in the NTD of the S protein is considered to be associated with antibody escape^12,13^. The Delta variant has been shown to have a higher replication rate, transmissibility, viral load as well as immune evasion^14–16^. The recently reported coronavirus variant B.1.1.529, named “Omicron” by WHO, is ringing the alarm bells around the world as it has 32 mutations in the spike protein and this might help the virus escape immunity. From the GISAID database, these 32 conserved Spike mutations are A67V, Δ69-70, T95I, G142D/Δ143-145, Δ211/L212I, ins214EPE, G339D, S371L, S373P, S375F, K417N, N440K, G446S, S477N, T478K, E484A, Q493K, G496S, Q498R, N501Y, Y505H, T547K, D614G, H655Y, N679K, P681H, N764K, D796Y, N856K, Q954H, N969K, L981F. Besides these, the conserved non-Spike mutations are -NSP3 – K38R, V1069I, Δ1265/L1266I, A1892T; NSP4 – T492I; NSP5 – P132H; NSP6 – Δ105-107, A189V; NSP12 – P323L; NSP14 – I42V; E – T9I; M – D3G, Q19E, A63T; N – P13L, Δ31-33, R203K, G204R ^17^. Among these variations, N679K, P681H (adjacent to the furin cleavage site), N501Y (within receptor binding motif), D614G (Spike protein protomer) have been earlier reported in other variants and found the variants be more transmissible and allow the virus to readily bind to the host cell ACE2R^7,18,19^. The mutation P681H has also been reported earlier in Alpha, Mu, some Gamma, and B.1.1.318 variants. According to WHO update on 28 November 2021, the current information about the transmissibility, disease severity, reinfection, effectiveness of existing vaccines, tests and treatment is not clear. Efforts are being made to better assess Omicron.

In the present *in silico* study, we have analysed the effect of mutations on the structure and binding affinity of the RBD region of Omicron and Delta variants (AY.1, AY.2, & AY.3) with ACE2R and with five different monoclonal SARS-CoV-2 neutralising human antibodies, namely CR3022, B38, CB6, P2B-2F6, and REGN. The MD simulation done in this study provides insights into the structural variations. The docking analysis of the RBD region of Omicron and Delta variants (AY.1, AY.2, & AY.3) with the ACE2 receptor (ACE2R) and with selected antibodies showed differences in binding affinity when compared with the wild SARS-CoV-2 (original strain) spike-RBD region.

## 2. Methodology

### 2.1 Data sets

The crystal structure of different human neutralizing monoclonal antibodies CR3022 6W41 ^20^, B38 7BZ5^21^, CB6 7C01^22^, P2B-2F6 7BWJ ^23^,and REGN 6XDG^24^ were retrieved from PDB RCSB database. In addition, the crystal structure of the hACE2 receptor (PDB ID: 7A97) and S protein (7AD1) was also retrieved^25^.

### 2.2 Creation of mutant structure and preprocessing

The Swiss model was used to create the RBD mutants (Omicron, Delta AY.1, AY.2, and AY.3) ^26^. In the Swiss model, 7AD1 was used as a template for homology modelling of mutations. A Modrefiner was also employed to reduce the energy of the mutant structure^27^. PDB-Sum was used to evaluate the simulated structure ^28^.The structure of the spike glycoprotein was preprocessed by eliminating all nonstandard residues, including water molecules, and replacing them with hydrogen atoms using the Discovery studio programme^29^. In addition, the monomeric structure of the protein was examined for further research. By eliminating the spike glycoprotein chain from the complex and other nonstandard residues with the discovery studio, other antibodies-based complex structures were retrieved. The structure of the ACE2R was similarly constructed and preprocessed.

### 2.3 Prediction of Physicochemical parameters, secondary structure and superimposition of structures

The Psipred online server ^30^ predicted the physicochemical characteristics and secondary structure of Omicron and wild RBD as well as the protein disorderness of wild, Omicron, and Delta variants. By using multalign, a chimaera tool was used to superimpose wild and mutant RBD structures. The distance matrix of the wild and mutant structures was calculated by the superpose tool ^31^. The difference distance matrix algorithm was used to visually discover substantial differences between any two structures.

### 2.4 Docking analysis

The PatchDock server ^32,33^ was used to dock RBD mutant variants with specified targets (ACE2R and distinct five monoclonal antibody structures), with an RMSD of 4.0 and complex type as default. The geometric form complementarity score was used to conduct the docking investigation. A higher score suggests a stronger affinity for binding. The docking scores and interaction at the RBD areas determine the outcome of the results. LigPlot plus v2.2 was used to view protein-protein and antibody-protein interactions^34^. Antibody scripts under the antibody loop numbering scheme, i.e., the KABAT Scheme and the DIMPLOT script algorithm package integrated into LigPlot plus v2.2, were used to perform molecular interactions of antibodies and ACE2R with RBD variants.

### 2.5 Molecular Simulation Dynamics

GROMACS was used to investigate the molecular dynamics of wild-type and mutant RBD Spike variants. On the basis of protein dynamics, MD simulation was used to produce time-dependent conformational alterations and protein modifications. The GROMACS96 54a7 force field ^35^ was used for the MD simulation study. To cope with dissolvable water surrounding protein, spc216.gro was utilised as a none-lite equilibrated 3 point dissolvable water model in a dodecahedron. The RBD wild type structure and mutations (Omicron, Delta AY.1, AY.2, & AY.3) were electrically neutralised by adding Na+59 and Cl-62, Na+68 and Cl-75, Na+76 and Cl-80 ions, and Na+90 and Cl-95 ions, respectively. The salt content was kept constant at 0.15 mol/L in all of the systems. Water molecules added to the wild RBD structure, Omicron, Delta AY.1 & AY.2, and Delta AY.3 were 20104, 23191, 23167, and 31118, respectively. At this phase, we kept the protein in the middle, at least 1.0 nm from the case edges. To minimise the energy required in the following phase, we adopted the steepest descent approach. The framework is then equilibrated at 300 K temperature and 1 atm for 100 ps using the canonical ensemble (NVT) (constant number of particles, volume, and temperature) and the canonical ensemble (NPT) outfit (constant number of particles, pressure, and temperature). We extended the MD run time to 100 ns (RBD Wild type, Delta Variants) and 20 ns (Omicron) after finishing the equilibrium measure. Gromacs tools (gmxrms, rmsf, gyrate hbond, and sasa) were used to calculate root mean square deviation (RMSD), root mean square fluctuation (RMSF), gyrate for radius of gyration (Rg), H-bond (for intramolecular H-bonds), and solvent accessible surface area. The XMGRACE application was used to visualise the MD trajectory data^36^.

## 3. Results

### 3.1 Physicochemical parameters, secondary structure and superimposition of structures

The superimposition of the RBD region of the wild (original strain) and selected mutant variants suggests structural changes which are indicated in figure 1. Secondary structure prediction analysis of Omicron has shown many changes in the helix, strands, and coils etc. as compared to wild. The results are shown in figures 2 (A) and 2 (B). Also, the analysis suggests changes in S protein physiological properties like polarity and hydrophobicity at multiple positions, as shown in figures 2 (C) and (D). The RBD variant protein was determined to be intrinsically disordered as compared to wild (Figure 3). Plaxco and Gross (2001) argue that protein disorder is critical to understanding protein function and folding mechanisms. Protein dysfunction has also been linked to disorders induced by protein mis-folding and aggregation in biology ^37,38^.

**Figure 1.**
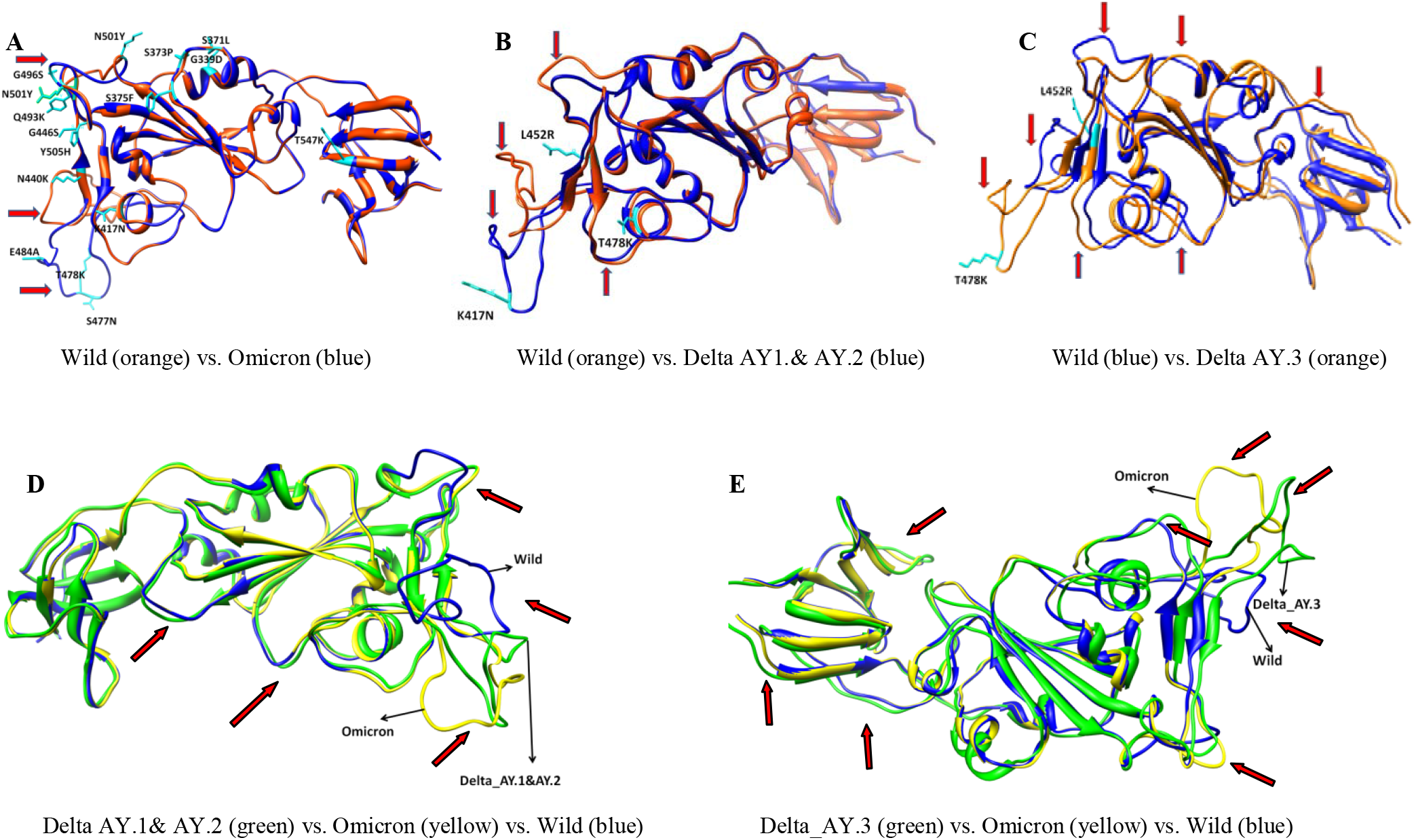
Structural superimposition of wild and mutant variants of the RBD region of Spike.Red arrows indicate the structurally hampered regions.

**Figure 2.**
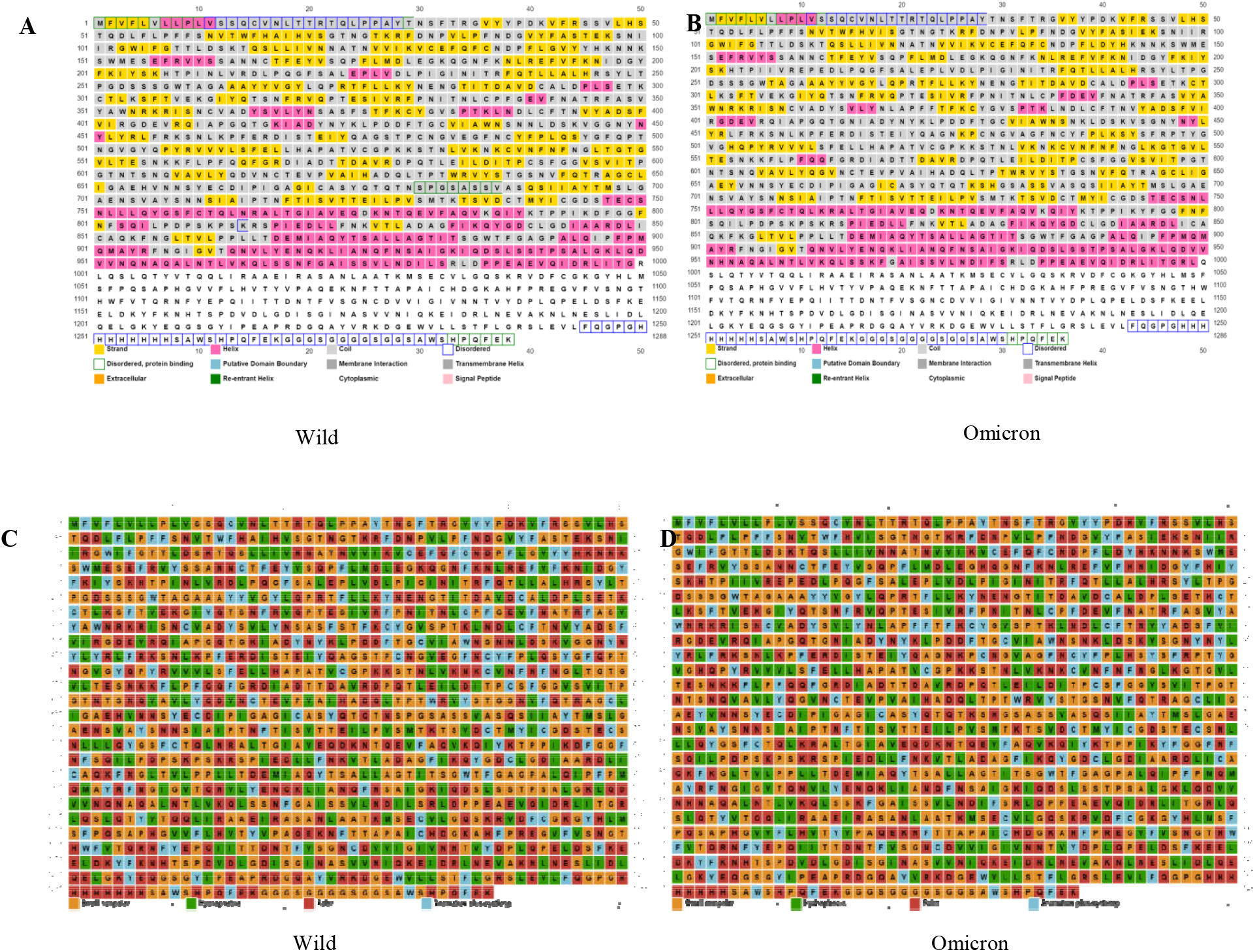
Prediction of secondary structure and physiochemical parameter changes in the wild and Omicron.

**Figure 3.**
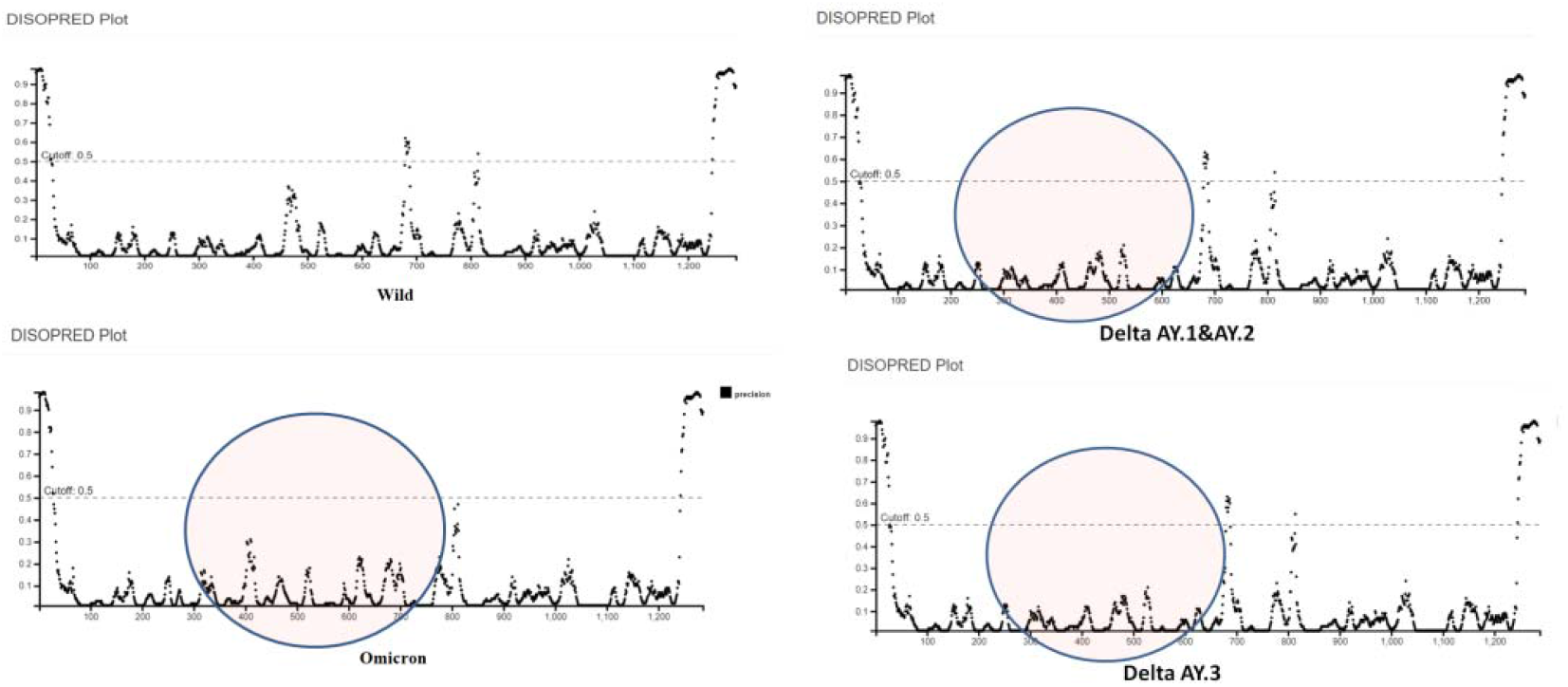
Prediction of protein disorderness.

All these results suggest that mutations may have altered the structures and physiochemical properties of Omicron as compared to wild and the Delta variants. The difference distance matrix results show that Delta variant AY.3 had a significant change in the overall RBD structure when compared to Delta AY.1 & AY.2 and Omicron (Figure 4).

**Figure 4.**
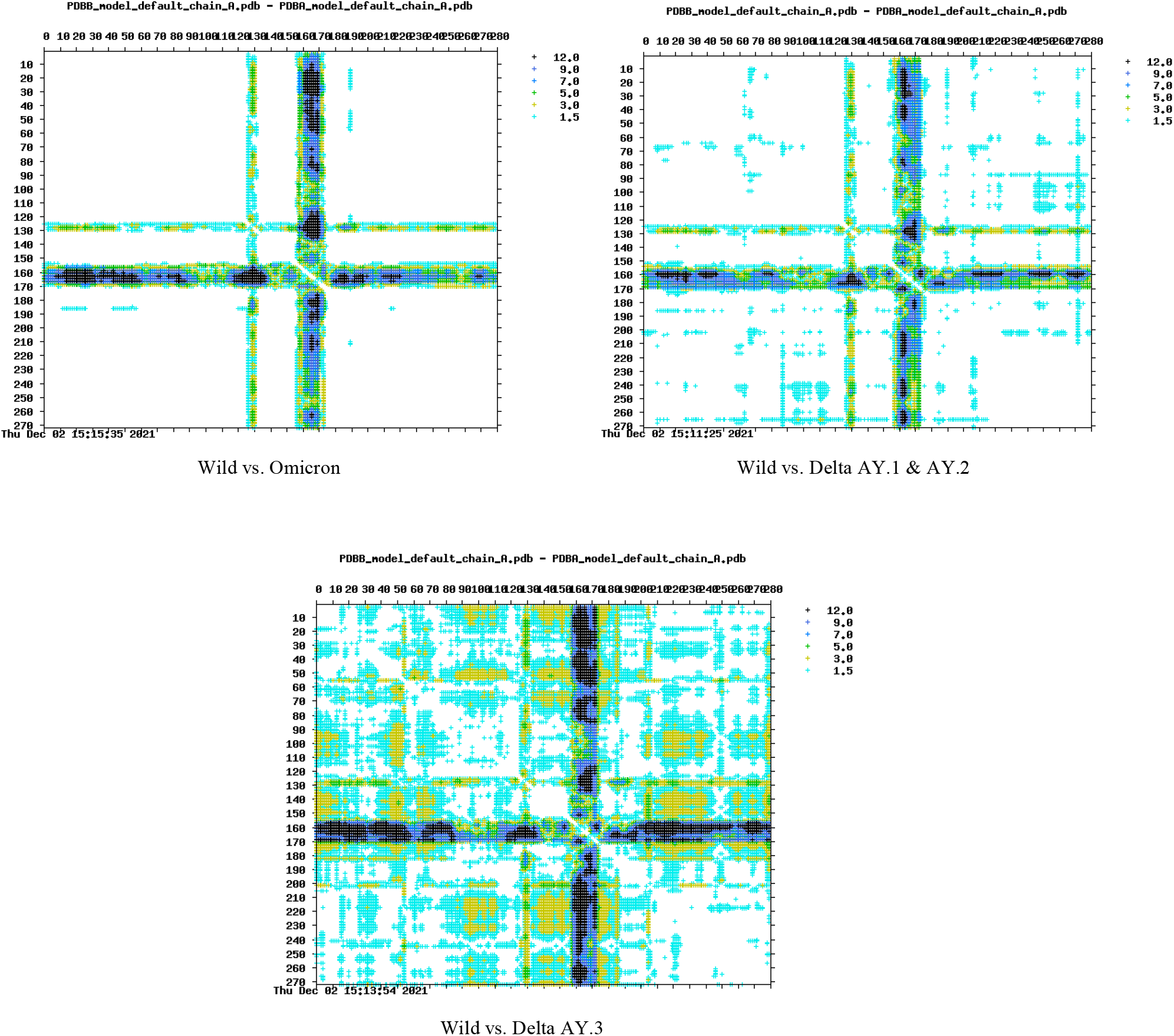
Prediction of RBD variants’ distance matrix (Wild, Omicron, Delta AY.1, AY.2 & AY.3).The lighter the region, the more similar are the structures. Likewise, the darker region corresponds to more prominent structures.Thedifference distance plot in Superpose shows six graded cutoffs. White depicts difference between 0 and 1.5 Angstroms, yellow depicts difference between 1.5 and 3.0 Angstroms, light green depictsdifference between 3.0 and 5.0 Angstroms, dark turquoise depictsdifference between 5 and 7 Angstroms, dark blue depictsdifference between 7 and 9 Angstroms, and black depicts difference between more than 9 Angstroms.

### 3.2 Docking analysis

The docking analysis of the RBD region of Omicron and Delta variants (AY.1, AY.2, & AY.3) with ACE2R showed differences in binding affinity when compared with the wild SARS-CoV-2 (original strain) spike-RBD region. The binding score determines the binding affinity. The higher the binding score, the higher the binding affinity. The binding score of ACE2R-Omicron RBD is higher (18208) than that of ACE2R-wild RBD, which is 17910, but it is less than ACE2R-Delta AY.3 (19084). The ACE2R-Delta AY.1 & 2 binding score 16886 is the lowest of all (Table 1).Through docking, the binding of the RBD region of Omicron with five different antibodies, viz. CR3022, B38, CB6, P2B-2F6, and REGN, was analysed and compared with wild SARS-CoV-2 (original strain) spike-RBD region and Delta variant sub-lineages (AY.1, AY.2, & AY.3). The binding scores are shown in table 2, and the different interactions are depicted in the supplementary file. The binding scores of antibodies CR3022, B38, CB6, P2B-2F6, and REGN with wild-RBD are 9248, 19152, 15984, 12776, and 14478, respectively. The binding score of antibodies CR3022, B38, CB6, P2B-2F6, and REGN with Omicron-RBD is 8768, 13240, 13660, 11900, and 14696, respectively, which is less than the binding score of wild-RBD binding except for REGN. Also, besides REGN, the binding score of the other four antibodies to Omicron-RBD is less than that of antibodies to Delta AY.1 and 2. The binding scores for Omicron-RBD vs. CR3022 (8768), Omicron-RBD vs. B38 (13240) and Omicron-RBD vs. P2B-2F6 (11900) are less than those of Delta AY. 3 sub-lineages (i.e., with CR3022 = 9158, B38 = 14308, and P2B-2F6 = 12124). The remaining two antibodies, CB6 and REGN, have a binding score slightly higher for Omicron-RBD (13660 and 14696) as compared to Delta AY. 3 sub-lineages (13206 and 13236).

**Table 1:** Table showing the binding score from docking of RBD region of each wild SARS-CoV-2 (original strain), Omicron and Delta variants (AY.1, AY.2, & AY.3) with ACE2 receptor (ACE2R)

| Receptor | Binding score of Spike-wild RBD | Binding score of Omicron | Binding affinity | Binding score of Delta AY.1 & 2 | Binding affinity | Binding score of Delta AY.3 | Binding affinity |
| --- | --- | --- | --- | --- | --- | --- | --- |
| ACE2R | 17910 | 18208 | Higher | 16886 | Lesser | 19084 | Higher |

**Table 2:** Table showing the binding score from docking of RBD region of each wild SARS-CoV-2 (original strain), Omicron and Delta variants (AY.1, AY.2, & AY.3) with CR3022, B38, CB6, P2B-2F6, and REGN antibodies.

| Antibodies | Binding score of Spike-wild RBD | Binding score of Omicron | Binding affinity | Binding score of Delta AY.1 & AY.2 | Binding affinity | Binding score of Delta AY.3 | Binding affinity |
| --- | --- | --- | --- | --- | --- | --- | --- |
| CR3022 | 9248 | 8768 | Lesser | 8878 | Lesser | 9158 | Lesser |
| B38 | 19152 | 13240 | Lesser | 15646 | Lesser | 14308 | Lesser |
| CB6 | 15984 | 13660 | Lesser | 14012 | Lesser | 13206 | Lesser |
| P2B-2F6 | 12776 | 11900 | Lesser | 13016 | Higher | 12124 | Lesser |
| REGN | 14478 | 14696 | Higher | 14122 | Lesser | 13236 | Lesser |

### 3.3 Molecular Simulations Dynamics

The RMSD values of wild-type and mutant proteins were compared to better understand the impact of mutations on protein structure. We calculated the RMSD for all proteins’ backbones with reference to their original structures during the molecular dynamics simulation. The structure of Omicron and Delta AY.3 swings more than wild and Delta AY.1 & AY.2 according to RMSD data (Figure 5). Wild, Omicron, Delta (AY.1 & AY.2), and Delta AY.3 had average RMSD values of 0.25 nm, 0.39 nm, 0.26 nm, and 0.43 nm, respectively. We also kept track of each atom’s RMSF variations to examine how the mutation changed the dynamic behaviour of the protein. Wild, Omicron, Delta (AY.1 & AY.2), and Delta AY.3 had average RMSF values of 0.16 nm, 0.17 nm, 0.15 nm, and 0.15 nm, respectively. In some regions, the RMSF value in Delta AY.3, Delta (AY.1 & AY.3) was greater (Figure 5). In Omicron and Delta, Rg fluctuation was greater (AY.1 & AY.2). In comparison to the wild, Delta AY.3 (Figure 5) showed the least changes. Wild, Omicron, Delta (AY.1 & AY.2), and Delta AY.3 have average Rg values of 1.979nm, 1.978 nm, 1.62 nm, and 1.67 nm, respectively. Wild, Omicron, Delta (AY.1 & AY.2), and Delta AY.3 have intramolecular h-bonding of 171.28, 174.04, 177.71, and 178.23, respectively (Figure 5). 158.48 nm2, 154.52 nm2, 154.31 nm2, and 152.34 nm2 were the SASA values in Wild, Omicron, Delta (AY.1 & AY.3), and Delta AY.3 correspondingly (Figure 5).

**Figure 5.**
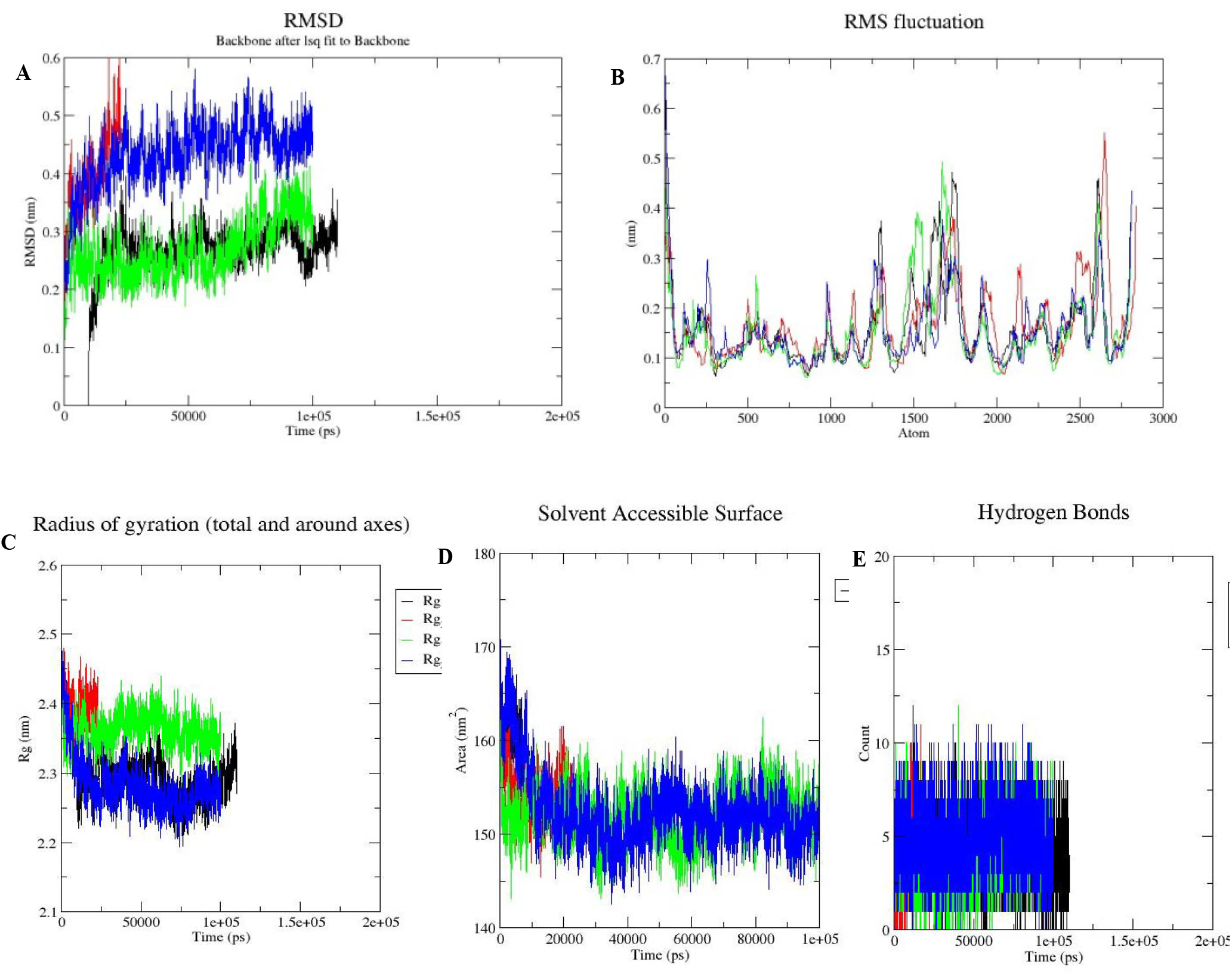
Molecular simulation results of wild type and mutant variants (Omicron, Delta AY.1 & AY.2, and Delta AY.3). (A) RMSD plots, (B) RMSF plots, and (C) radius of gyration (Rg) plots, (D) Solvent accessible surface area (SASA), intramolecular H-bonds (E).Wild type□black, Omicron□red, Delta AY.1 & AY.2- Green, and Delta AY.3- Blue.

## 4. Discussion

Genetic lineages of SARS-CoV-2 have been arising and spreading around the world since the commencement of the COVID-19 pandemic. The S protein mediates the attachment of the virus to the host cell-surface receptor that making it a major target of neutralizing antibodies following SARS-CoV-2 infection ^39,40^. SARS-CoV-2 S Protein has a major role in host-pathogen interaction^41^. As per the WHO report and its international networks of experts till now a number of mutations in genomes of SARS-CoV-2 virions have been reported that are expected to be either neutral or moderately detrimental. Some mutations are expected to impact virus biology by affecting viral antigenicity, transmissibility, pathogenicity, and infectivity. Mutations in the spike protein of SARS-CoV-2 are expected to alter the antigenic phenotype of SARS-CoV-2 and affect immune recognition too that requires immediate attention ^42,43^. UK variant of B.1.1.7 lineage has 8 mutations in S protein that seems to be remaining susceptible for RNA-based COVID-19 vaccine BNT162b2 ^44^. South African variant of lineage B.1.351 having E484K mutations reported being incompliant for neutralizing antibody as well as convalescent plasma and sera from the vaccinated population ^45^. A variant of B.1.1.28 lineage first identified in Brazil has 10 RBD mutations and one within the furin cleavage site. This variant was also found to have resistance for RBD targeted neutralizing antibody similar to B.1.351 ^46,47^. Lineage B.1.526 that contains E484K variation has been reported in New York first and found to have resistance for therapeutic monoclonal antibodies meanwhile low susceptibility for neutralization by vaccine sera or convalescent plasma ^48^. In 2021 B.1.67 (L452R & E484Q) RBD variant was reported in India which brought a deadlier second wave in the country within a short time. *In silico* analysis of structural stability and molecular simulation data of double mutant predicted reduced binding affinity for CR3022 antibody as well as lower vaccine efficacy of double mutant with antibody in comparison to wild type ^35^. The B.1.617.2 lineage named as delta variant has become a more transmissible and dominant strain and the structural changes due to mutation may have caused the reduced response to vaccines. A study has shown that the mutation in the Delta variant causes reduced binding with neutralizing antibodies and thereby escaping the immunity ^49^.

The newly emerged omicron variant has been found to have the highest number of variations among all earlier reported SARS-CoV-2 variants (32 RBD variations and 18 non-RBD variations) which invoke an evaluation of the potential of infectivity and contagious characteristics of this variant as well as vaccine and antibody efficacy. Here we performed computational analysis of the changed structure of spike glycoprotein of Omicron variant and also compared with most transmissible and dominant Delta variants (AY.1, AY.2, & AY.3) as well as wild SARS-CoV-2 (original strain) to investigate the discrepancy in susceptibility to the infection and potential for immune evasion. To estimate the binding efficiency of the variant’s RBD region to ACE2R we carried out docking analysis of the RBD region to ACE2R. The binding score of ACE2R-Omicron RBD is 18208 while for ACE2R-wild RBD is 17910, ACE2R-Delta AY.3 is 19084 and ACE2R-Delta AY.1 & 2 are 16886 which shows that the omicron’s RBD has higher binding affinity for ACE2R compared to wild SARS-CoV-2 original strain and Delta AY.1 & 2 whereas lower affinity compared to Delta AY.3 strain. The binding affinity of Delta AY.1 & 2 for ACE2R is least among all. Docking for evaluating binding efficiency of RBD region for five different humanmonoclonal antibodies viz. CR3022, B38, CB6, P2B-2F6, and REGN has also been done and found that the binding score of Omicron-RBD for antibodies CR3022, B38, CB6, P2B-2F6, and REGN is 8768, 13240, 13660, 11900, and 14696 respectively which is lesser than the binding score of wild-RBD as well as Delta AY.1 & 2 except for REGN. The binding score for Omicron-RBD for CR3022, B38, and P2B-2F6 is lesser than that of Delta AY.3 sub-lineage while for CB6 and REGN binding score is slightly higher for Omicron-RBD as compared to Delta AY.3 sub-lineage. Our docking analysis revealed lower interaction of Omicrons-RBD with human antibodies compared to wild strain of SARS-CoV-2 and deadly Delta variant and higher interaction with ACE2R compared to wild strain of SARS-CoV-2 and Delta AY.1 & 2 while lower in comparison to Delta AY.3. Next, we superimposed the 3D structure of the RBD region of SARS-CoV-2 wild strain with Omicron, Delta AY.1, AY.2 & AY.3 to predict alteration in structural and physiochemical parameters. This analysis shows intrinsically disordered Omicron-RBD protein along with a change in helix, strand, and the coil of Omicron structure as compared to the wild and Delta variant strains. Besides structural changes, polarity and hydrophobicity of Omicron were also found different from the wild type which indicates that the omicron variant is structurally and physiochemicaly different from wild stain however major alterations in whole RBD structure were found in Delta variants AY.3 in contrast to Delta AY.1 & AY.2 and Omicron. MD simulation was done to analyze the impact of mutations on protein structure. MD simulations provide insights into protein’s behavior in its natural environment and compute its trajectory over time, providing information on changed protein structure and fluctuations that may help to analyze flexibility and stability ^50^. Pathological phenotypes can be caused by changes in protein stability and flexibility ^51^. We evaluate the transient characteristics of wild-type and mutant Spike RBD to understand the functional and structural differences between the two. High fluctuations of RMSD were observed in Omicron and Delta AY.3 variants as compared with wild type while lower in Delta AY.1 & AY.2. The RMSD analysis showed lower average RMSD value of wild type in order [wild type < Delta (AY.1 & AY.2) < Omicron < Delta AY.3] suggesting wild RBD was more stabilized and Delta AY.3 and Omicron showed unstable structure. A higher RMSF value was observed in Omicron that shows a more flexible structure than other variants which showed limited movement in the structure. In comparison to wild, the lowest Rg value found in Delta (AY.1 &AY.2) followed by Delta AY.3 and Omicron that intimates more compactness in protein structure. Intramolecular HLbonds showed higher fluctuations in all mutant variants compared with wild type; albeit they overlapped at times and were higher and lower than wild type at other periods in the case of Delta (AY.1 & AY.2), Delta AY.3, and Omicron (Figure 5). Intramolecular HLbonding showed that the wild type had a lower average number of hydrogen bonds than Omicron, Delta (AY.1 & AY.2), and Delta AY.3 sequentially. The stiffness of proteins and their interactions may be affected by fluctuations in total intramolecular HLbonds ^52^. The highest SASA value was observed in the wild type than the variants in decreasing order: wild > Omicron >Delta (AY.1 & AY.3) > Delta AY.3 which indicate that the examined variants are less accessible than the wild-type protein, which might affect their capacity to interact with other molecules.

## 5. Conclusion

In summary this study anticipated more binding affinity of Omicron variant with ACE2R while lower affinity for neutralizing antibodies in contrast to wild type and Delta variant AY.1 & 2. However, Delta AY.3 shows highest binding affinity for ACE2R in contrast to Omicron variant. In addition Delta AY.3 and Omicron variant is likely to be relatively unstable and highly compact protein structure that may lead to more pathogenicity as well as antibody escape than wild and Delta AY.1 & 2 strains. According to MDand docking analysis, Delta variant AY.3 RBD region is majorly affected in comparison to Delta AY.1 & AY.2 and Omicron and shows higher binding with ACE2R, unstable structure, hampered interactions with antibodies and thus could still believed to be more pathogenic and immune evasive.Omicron had reduced binding effectiveness for CR3022, B38, CB6, and P2B2F6 than wild and Delta AY.1 and AY.2 according to the data. However, as compared to Delta AY.3, the Omicron RBD region had a reduced binding effectiveness for CR3022, B38, and P2B2F6 antibodies, which might lead to antibody escape.The Delta variants (AY.1, AY.2, AY.3) have D614G, P681R, D950N changes in the fusion region that binds to host receptor while Omicron has more changes (D614G, H655Y, N679K, P681H, N764K, D796Y, N856K, Q954H) in the fusion region. This may affect the host-pathogen interactions and ultimately transmissibility which need further validation from real-world data.

## Abbreviations

H□bonding: hydrogen bonding
Rg: radius of gyration
RMSD: root mean square deviations
RMSF: root mean square fluctuations
SASA: Solvent Accessible Surface Area

## 6. Acknowledgements

Banaras Hindu University in Varanasi, India, supplied the internet and computer resources for this project. Prashant Ranjan, Neha are supported by the RET fellowship at BHU; Chandra Devi, DBT JRF; Kaviyapriya A. D Summer intern supported by Indian Academy of Sciences Fellowship.

## 7. Competing interests

The authors declare that they have no any conflict of interest

## 8. Author Contributions

PR: Conceptualization, literature mining, bioinformatics analysis, MD simulations, data analysis and manuscript writing;Neha: Literature mining, manuscript writing, and Data analysis; CD: Literature mining, manuscript writing, and Data analysis. K.A.D: Data analysis PD: Major inputs, correspondence, corrections in the manuscript.

## Notes

### Competing Interest Statement

The authors have declared no competing interest.

## References

1. Fleischmann Jr, W. R. Viral genetics. Med. Microbiol. 4th Ed. (1996).

2. Huang, Y. et al. Clinical characteristics of laboratory confirmed positive cases of SARS-CoV-2 infection in Wuhan, China: A retrospective single center analysis. Travel Med. Infect. Dis. 36, 101606 (2020).

3. Lopez Bernal, J. et al. Effectiveness of Covid-19 vaccines against the B. 1.617. 2 (Delta) variant. N Engl J Med 585–594 (2021).

4. Fehr, A. R. & Perlman, S. Coronaviruses: an overview of their replication and pathogenesis. Coronaviruses 1–23 (2015).

5. Hoffmann, M. et al. SARS-CoV-2 cell entry depends on ACE2 and TMPRSS2 and is blocked by a clinically proven protease inhibitor. Cell 181, 271–280 (2020).

6. Huang, Y., Yang, C., Xu, X., Xu, W. & Liu, S. Structural and functional properties of SARS-CoV-2 spike protein: potential antivirus drug development for COVID-19. Acta Pharmacol. Sin. 41, 1141–1149 (2020).

7. Zhang, L. et al. SARS-CoV-2 spike-protein D614G mutation increases virion spike density and infectivity. Nat. Commun. 11, 1–9 (2020).

8. Ahamad, S., Kanipakam, H. & Gupta, D. Insights into the structural and dynamical changes of spike glycoprotein mutations associated with SARS-CoV-2 host receptor binding. J. Biomol. Struct. Dyn. 1–13 (2020).

9. Fuentes-Prior, P. Priming of SARS-CoV-2 S protein by several membrane-bound serine proteinases could explain enhanced viral infectivity and systemic COVID-19 infection. J. Biol. Chem. 296, (2021).

10. Korber, B. et al. Tracking changes in SARS-CoV-2 Spike: evidence that D614G increases infectivity of the COVID-19 virus. Cell 182, 812–827 (2020).

11. Bian, L. et al. Impact of the Delta variant on vaccine efficacy and response strategies. Expert Rev. Vaccines 20, 1201–1209 (2021).

12. Baj, A. et al. Spike protein evolution in the SARS-CoV-2 Delta variant of concern: a case series from Northern Lombardy. Emerg. Microbes Infect. 1–10 (2021).

13. Chaudhari, A., Kumar, D., Joshi, M. & Patel, A. E156/G and Arg158, Phe-157/del mutation in NTD of spike protein in B. 1.167. 2 lineage of SARS-CoV-2 leads to immune evasion through antibody escape. bioRxiv (2021).

14. Bolze, A. et al. Rapid displacement of SARS-CoV-2 variant B. 1.1. 7 by B. 1.617. 2 and P. 1 in the United States. medRxiv (2021).

15. Li, B. et al. Viral infection and transmission in a large well-traced outbreak caused by the Delta SARS-CoV-2 variant. MedRxiv (2021).

16. Mlcochova, P. et al. SARS-CoV-2 B. 1.617. 2 Delta variant replication and immune evasion. Nature 1–6 (2021).

17. England, P. H. SARS-CoV-2 variants of concern and variants under investigation in England. Tech. Brief. 30 (2021).

18. Scheepers, C. et al. Emergence and phenotypic characterization of C. 1.2, a globally detected lineage that rapidly accumulated mutations of concern. medRxiv 2008–2021 (2021).

19. Tao, K. et al. The biological and clinical significance of emerging SARS-CoV-2 variants. Nat. Rev. Genet. 1–17 (2021).

20. Yuan, M. et al. A highly conserved cryptic epitope in the receptor binding domains of SARS-CoV-2 and SARS-CoV. Science (80-.). 368, 630–633 (2020).

21. Wu, Y. et al. A noncompeting pair of human neutralizing antibodies block COVID-19 virus binding to its receptor ACE2. Science (80-.). 368, 1274–1278 (2020).

22. Shi, R. et al. A human neutralizing antibody targets the receptor-binding site of SARS-CoV-2. Nature 584, 120–124 (2020).

23. Ju, B. et al. Human neutralizing antibodies elicited by SARS-CoV-2 infection. Nature 584, 115–119 (2020).

24. Hansen, J. et al. Studies in humanized mice and convalescent humans yield a SARS-CoV-2 antibody cocktail. Science (80-.). 369, 1010–1014 (2020).

25. Fatihi, S. et al. A rigorous framework for detecting SARS-CoV-2 spike protein mutational ensemble from genomic and structural features. bioRxiv (2021).

26. Lyskov, S. & Gray, J. J. The RosettaDock server for local protein–protein docking. Nucleic Acids Res. 36, W233–W238 (2008).

27. Xu, D. & Zhang, Y. Improving the physical realism and structural accuracy of protein models by a two-step atomic-level energy minimization. Biophys. J. 101, 2525–2534 (2011).

28. Laskowski, R. A., Jabłońska, J., Pravda, L., Vařeková, R.S. & Thornton, J. M. PDBsum: Structural summaries of PDB entries. Protein Sci. 27, 129–134 (2018).

29. Systèmes, D. Biovia, discovery studio modeling environment. Dassault Systèmes Biovia San Diego, CA, USA (2016).

30. McGuffin, L. J., Bryson, K. & Jones, D. T. The PSIPRED protein structure prediction server. Bioinformatics 16, 404–405 (2000).

31. Maiti, R., Van Domselaar, G. H., Zhang, H. & Wishart, D. S. SuperPose: a simple server for sophisticated structural superposition. Nucleic Acids Res. 32, W590–W594 (2004).

32. Ranjan, P. & Das, P. Understanding the impact of missense mutations on the structure and function of the EDA gene in XLlinked hypohidrotic ectodermal dysplasia: A bioinformatics approach. J. Cell. Biochem. (2021).

33. Ranjan, P., Mohapatra, B. & Das, P. A rational drug designing: What bioinformatics approach tells about the wisdom of practicing traditional medicines for screening the potential of Ayurvedic and natural compounds for their inhibitory effect against COVID-19 Spike, Indian strain Spike, Papa. (2020).

34. Wallace, A. C., Laskowski, R. A. & Thornton, J. M. LIGPLOT: a program to generate schematic diagrams of protein-ligand interactions. Protein Eng. Des. Sel. 8, 127–134 (1995).

35. Ranjan, P., Neha Devi, C. & Das, P. Bioinformatics analysis of SARS-CoV-2 RBD mutant variants and insights into antibody and ACE2 receptor binding. bioRxiv 2021.04.03.438113 (2021) doi:10.1101/2021.04.03.438113.

36. Srikumar, P. S., Rohini, K. & Rajesh, P. K. Molecular dynamics simulations and principal component analysis on human laforin mutation W32G and W32G/K87A. Protein J. 33, 289–295 (2014).

37. Bates, S. H. et al. STAT3 signalling is required for leptin regulation of energy balance but not reproduction. Nature 421, 856–859 (2003).

38. Schweers, O., Schönbrunn-Hanebeck, E., Marx, A. & Mandelkow, E. Structural studies of tau protein and Alzheimer paired helical filaments show no evidence for beta-structure. J. Biol. Chem. 269, 24290–24297 (1994).

39. Piccoli, L. et al. Mapping neutralizing and immunodominant sites on the SARS-CoV-2 spike receptor-binding domain by structure-guided high-resolution serology. Cell 183, 1024–1042 (2020).

40. Liu, L. et al. Potent neutralizing antibodies against multiple epitopes on SARS-CoV-2 spike. Nature 584, 450–456 (2020).

41. Yan, R. et al. Structural basis for the recognition of SARS-CoV-2 by full-length human ACE2. Science (80-.). 367, 1444–1448 (2020).

42. Li, Q. et al. The impact of mutations in SARS-CoV-2 spike on viral infectivity and antigenicity. Cell 182, 1284–1294 (2020).

43. Letko, M., Marzi, A. & Munster, V. Functional assessment of cell entry and receptor usage for SARS-CoV-2 and other lineage B betacoronaviruses. Nat. Microbiol. 5, 562–569 (2020).

44. Muik, A. et al. Neutralization of SARS-CoV-2 lineage B. 1.1. 7 pseudovirus by BNT162b2 vaccine–elicited human sera. Science (80-.). 371, 1152–1153 (2021).

45. Wang, P. et al. Antibody resistance of SARS-CoV-2 variants B. 1.351 and B. 1.1. 7. Nature 593, 130–135 (2021).

46. Faria, N. R. et al. Genomic characterisation of an emergent SARS-CoV-2 lineage in Manaus: preliminary findings. Virological 372, 815–821 (2021).

47. Harvey, W. T. et al. SARS-CoV-2 variants, spike mutations and immune escape. Nat. Rev. Microbiol. 19, 409–424 (2021).

48. Annavajhala, M. K. et al. Emergence and expansion of the SARS-CoV-2 variant B. 1.526 identified in New York. medRxiv 2002–2021 (2021).

49. Baral, P. et al. Mutation-induced changes in the receptor-binding interface of the SARS-CoV-2 Delta variant B. 1.617. 2 and implications for immune evasion. Biochem. Biophys. Res. Commun. 574, 14–19 (2021).

50. Krebs, B. B. & De Mesquita, J. F. Amyotrophic lateral sclerosis type 20-In Silico analysis and molecular dynamics simulation of hnRNPA1. PLoS One 11, e0158939 (2016).

51. Khan, F. I., Wei, D.-Q., Gu, K.-R., Hassan, M. I. & Tabrez, S. Current updates on computer aided protein modeling and designing. Int. J. Biol. Macromol. 85, 48–62 (2016).

52. Hubbard, R. E. & Haider, M. K. Hydrogen bonds in proteins: role and strength. eLS (2010).

