## Supplemental File1 for "The influence of new SARS-CoV-2 variant Omicron (B.1.1.529) on vaccine efficacy, its correlation to Delta Variants: a computational approach"

**Supplementary File1**

Molecular interactions of five different antibodies, including CR3022, B38, CB6, P2B-2F6, and REGN, with structurally altered RBD variants, including Omicron, Delta, AY.1, AY.2, and AY.3 RBD. Hydrogen bonds are depicted in green dashed lines, and hydrophobic contacts are indicated by the red brick.

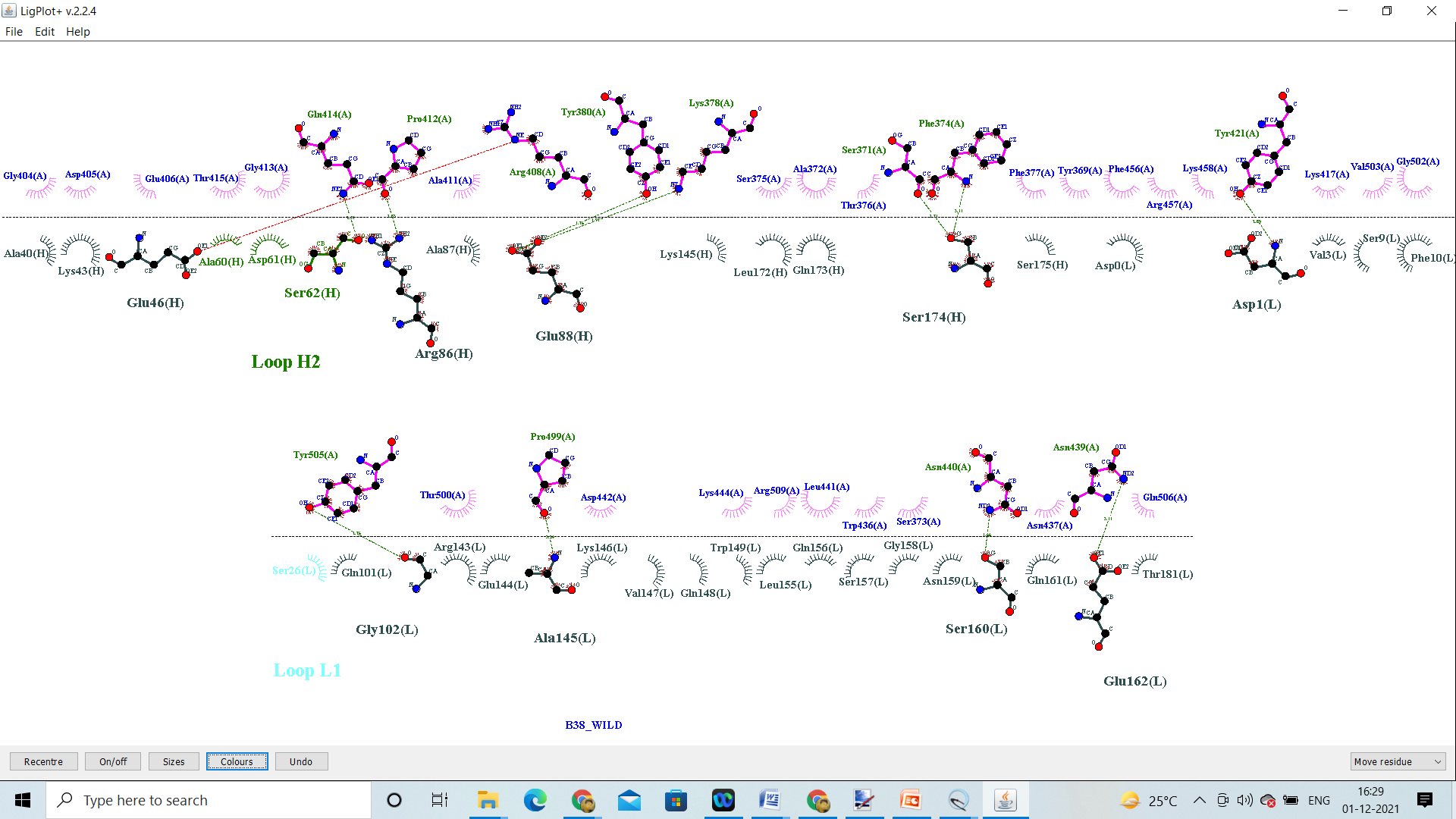

B38_Wild

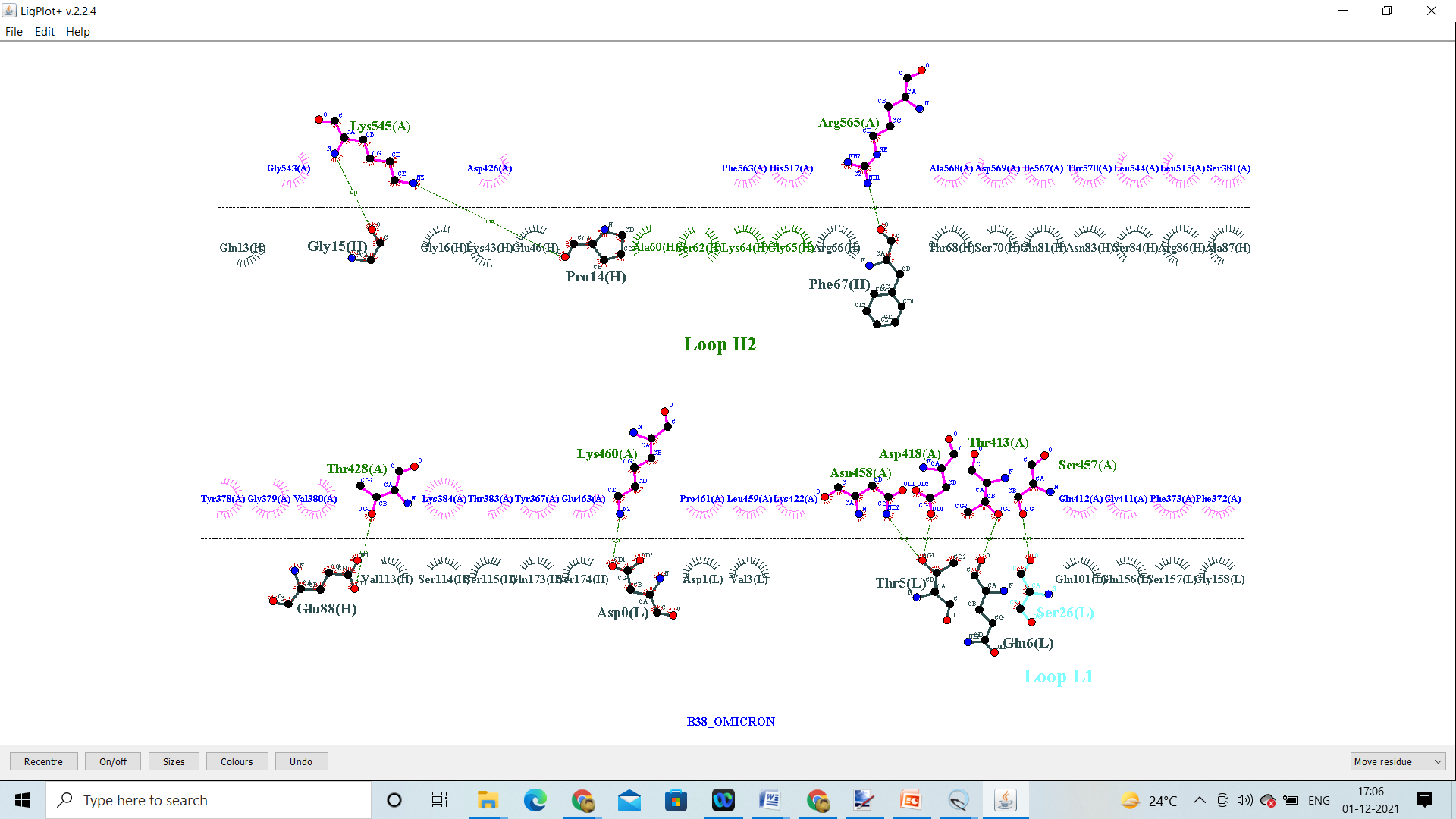

Q498Y

R408I

B38_Omicron

L455Y

K417G

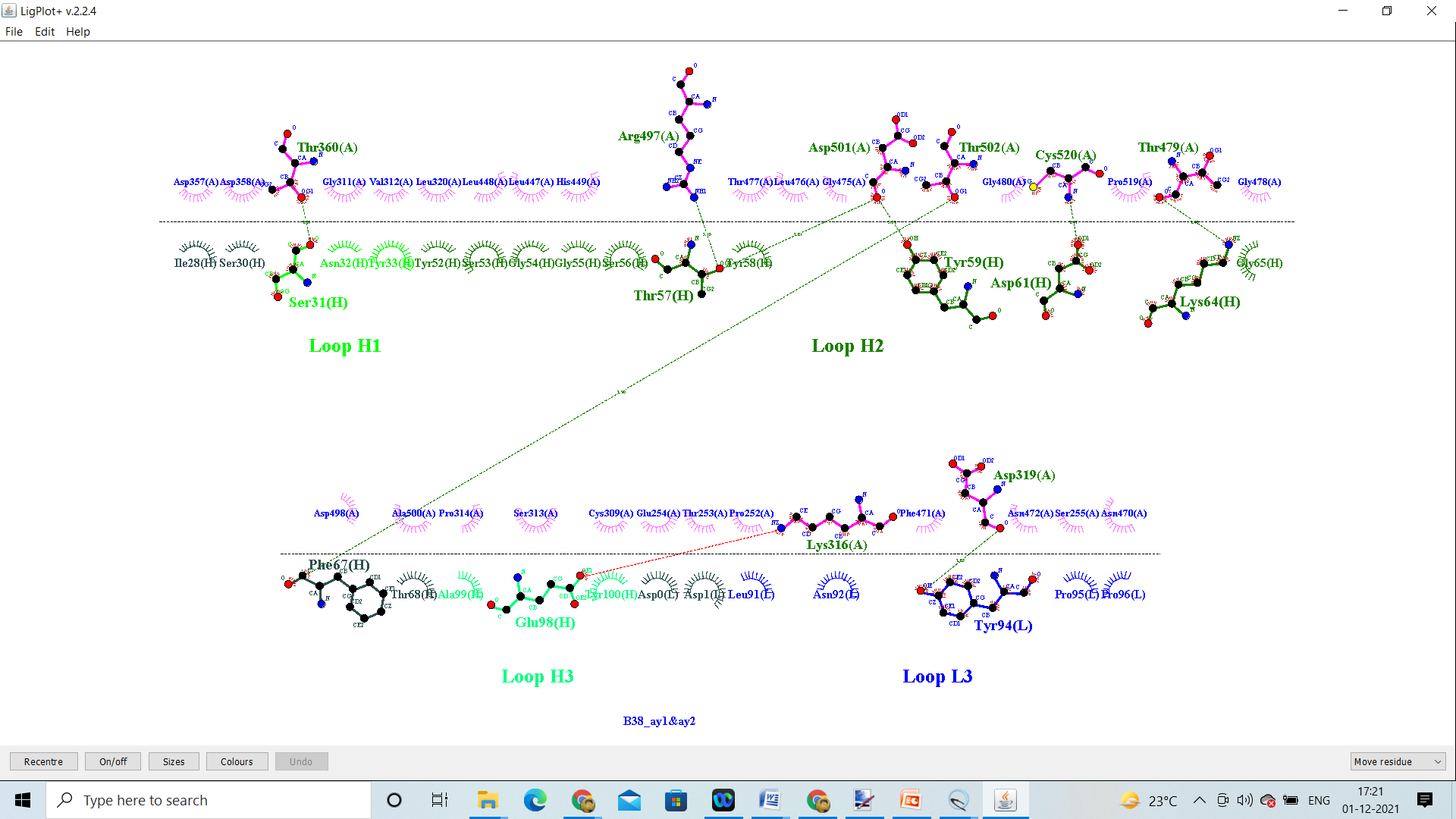

B38_Delta_AY.1&AY.2

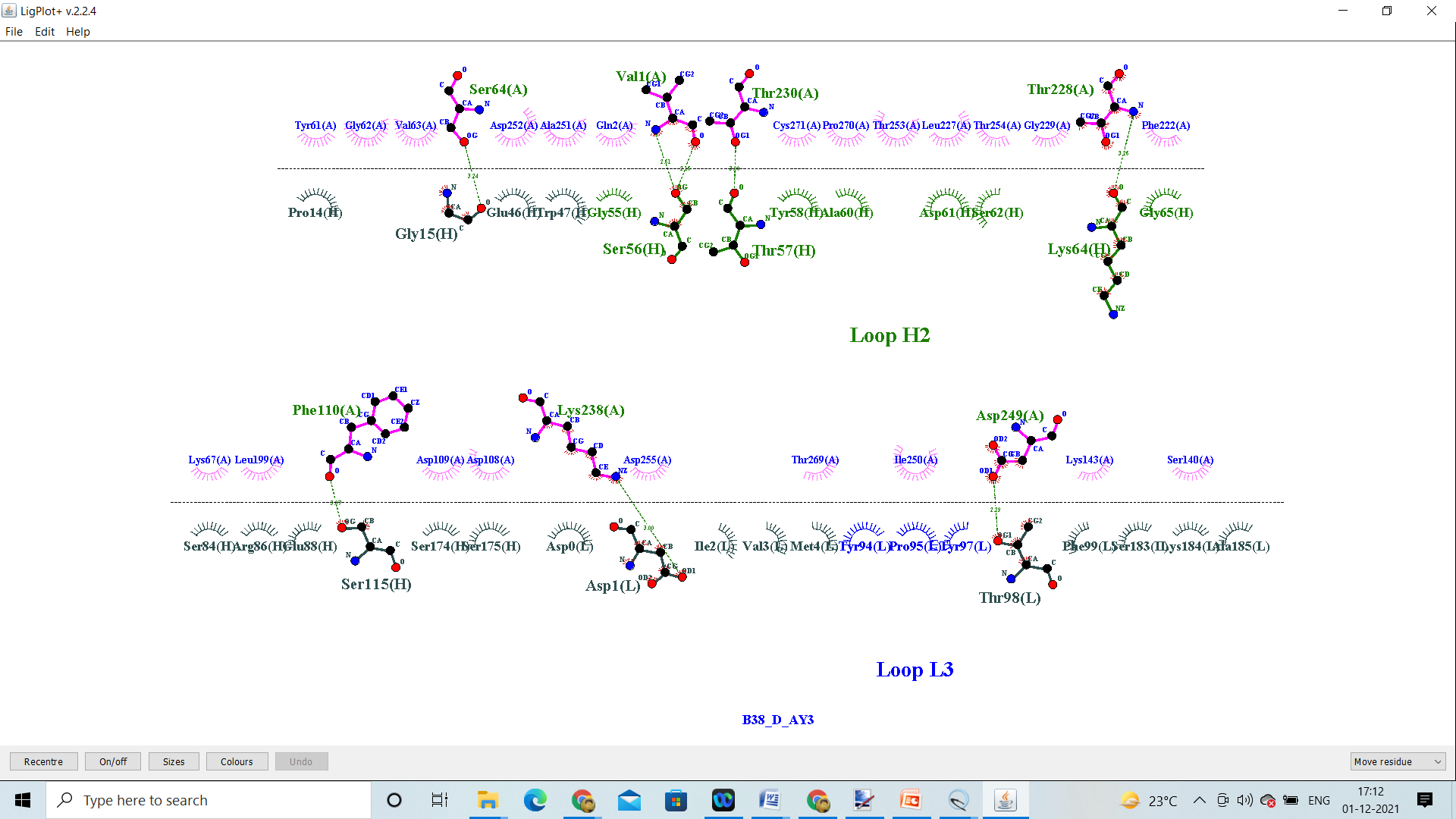

B38_Delta_AY.3

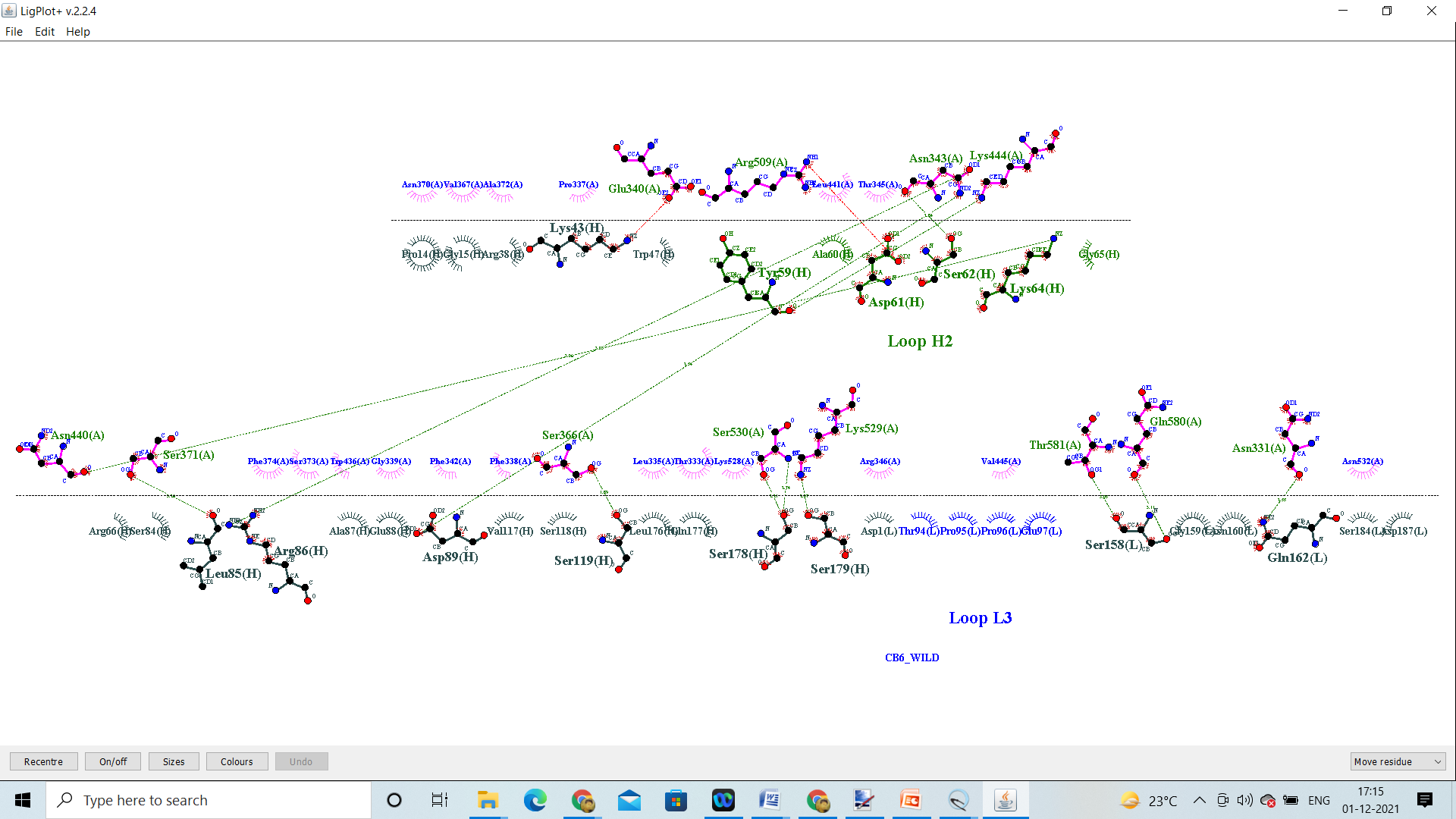

CB6_Wild

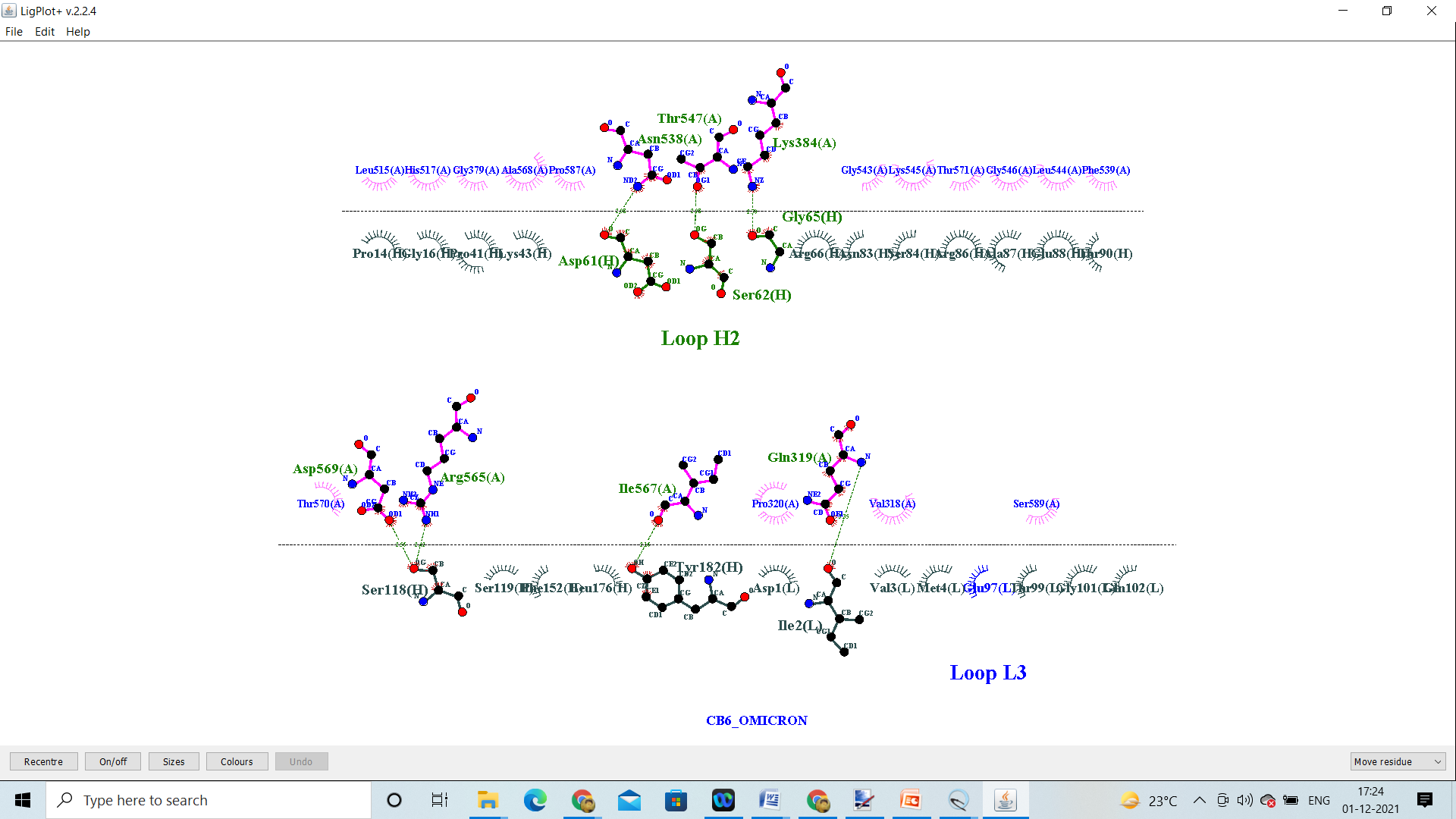

CB6_Omicron

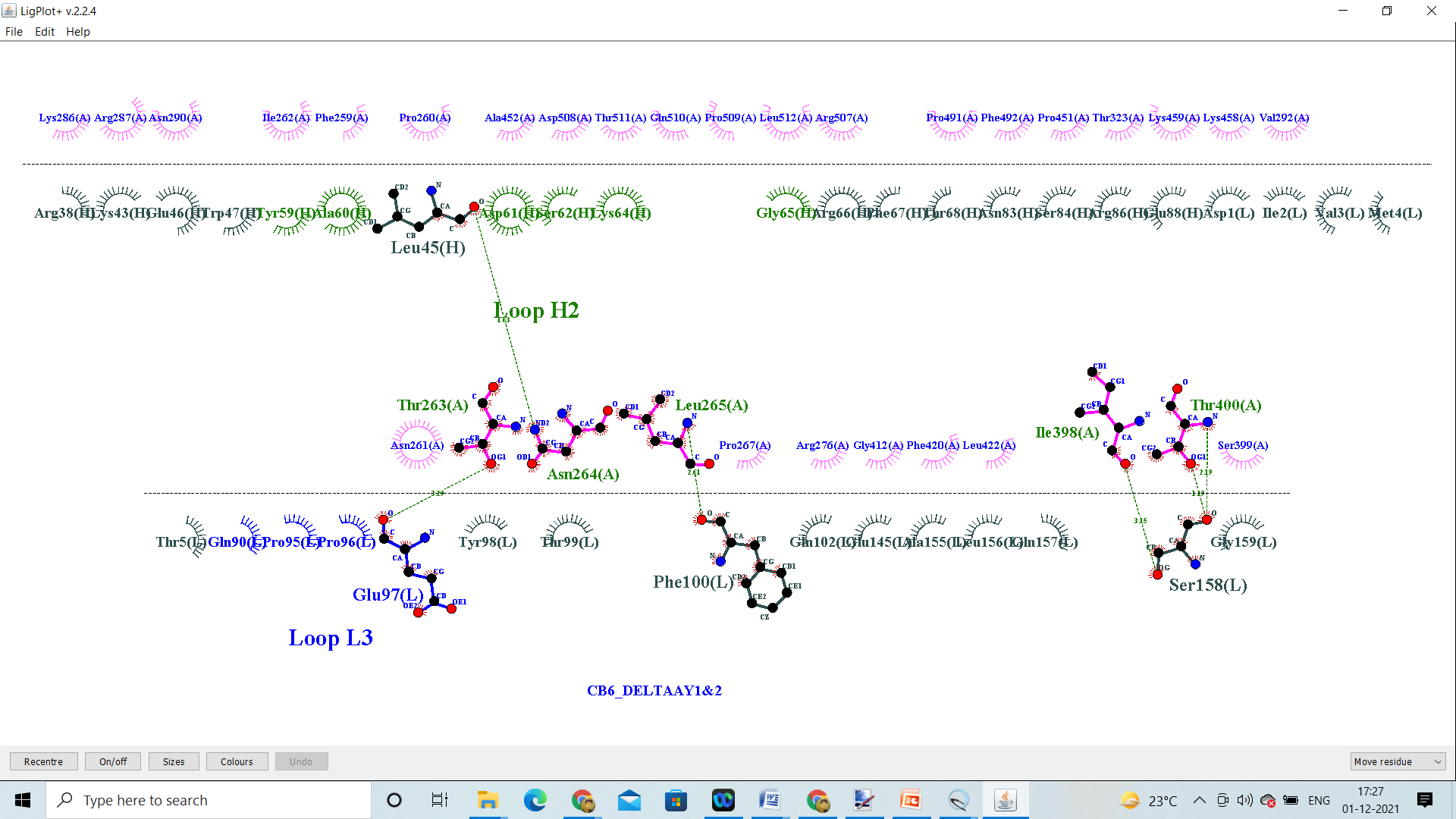

B38_Delta_AY.3

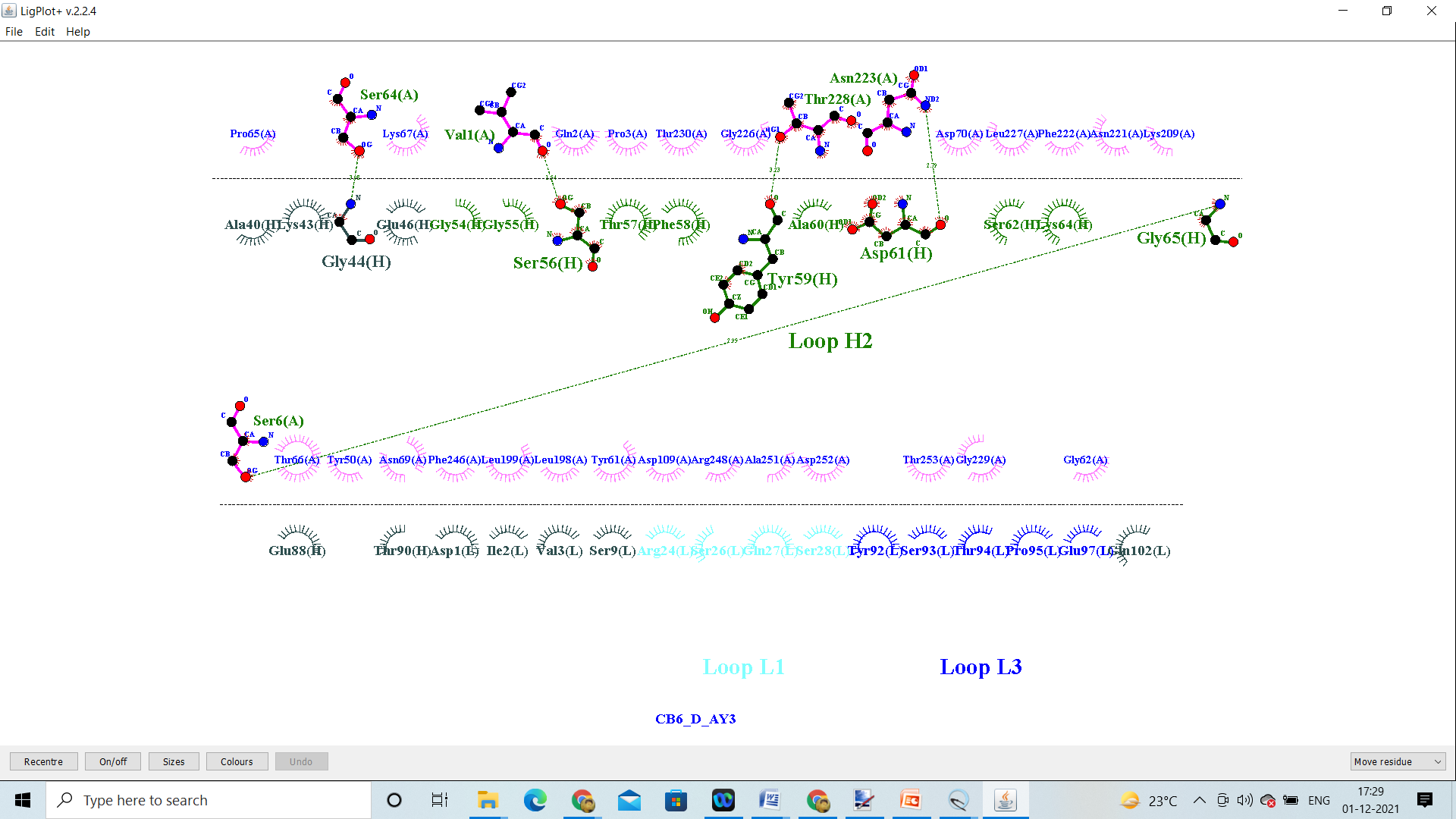

CB6_D_AY.3

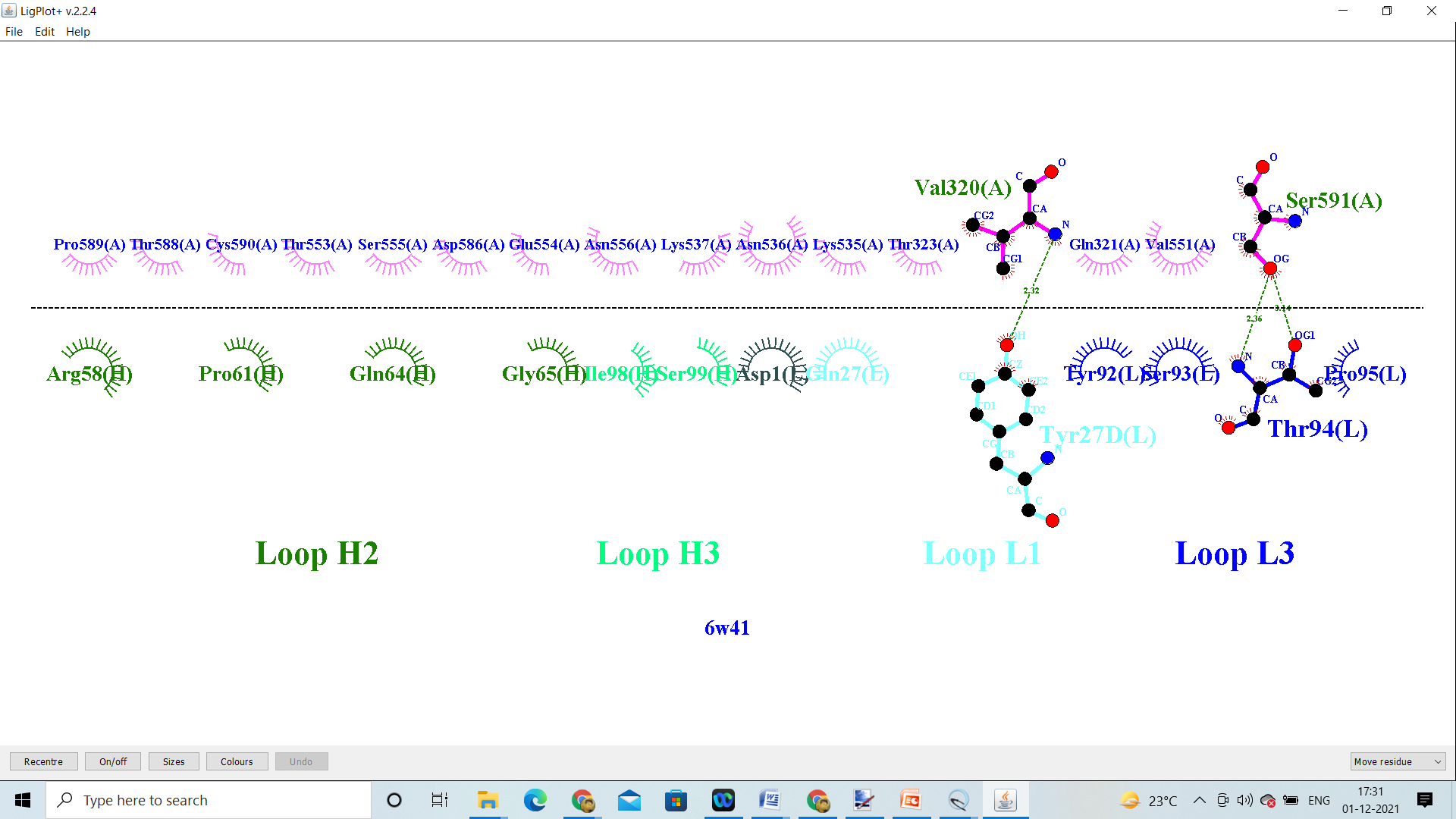

CR3022_Wild

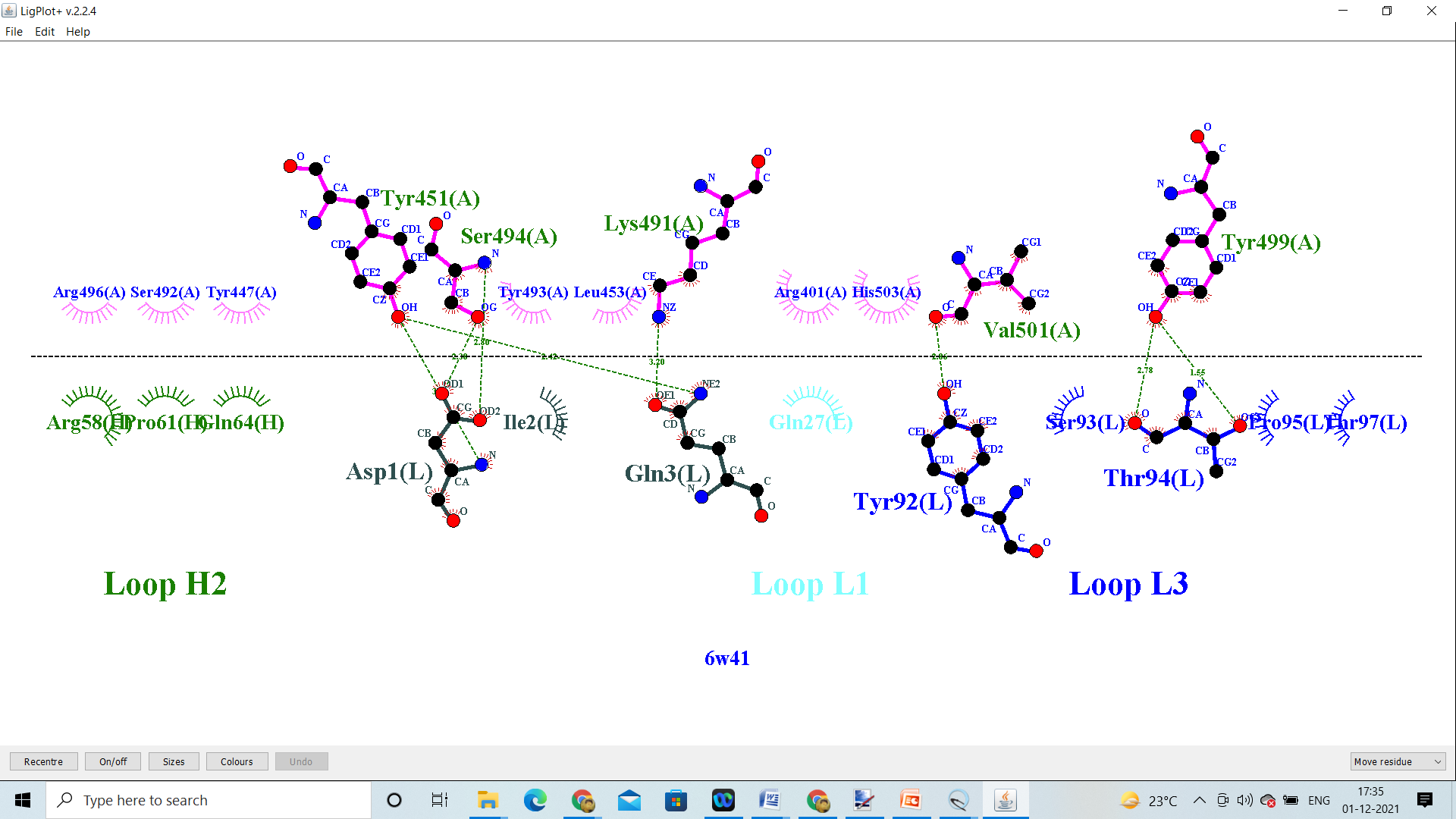

CR3022_Omicron

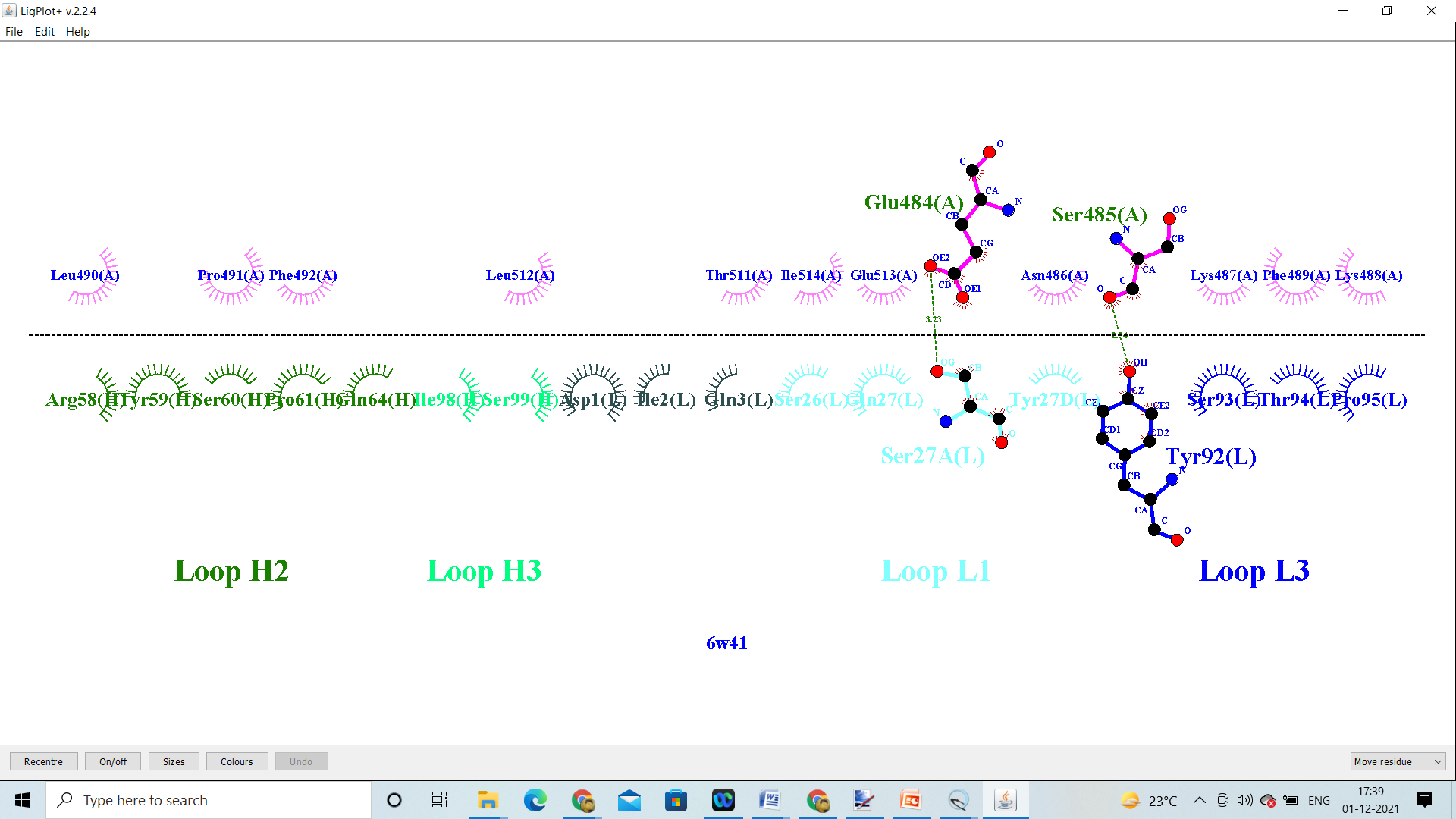

CR3022_Delta AY.1 &AY.2

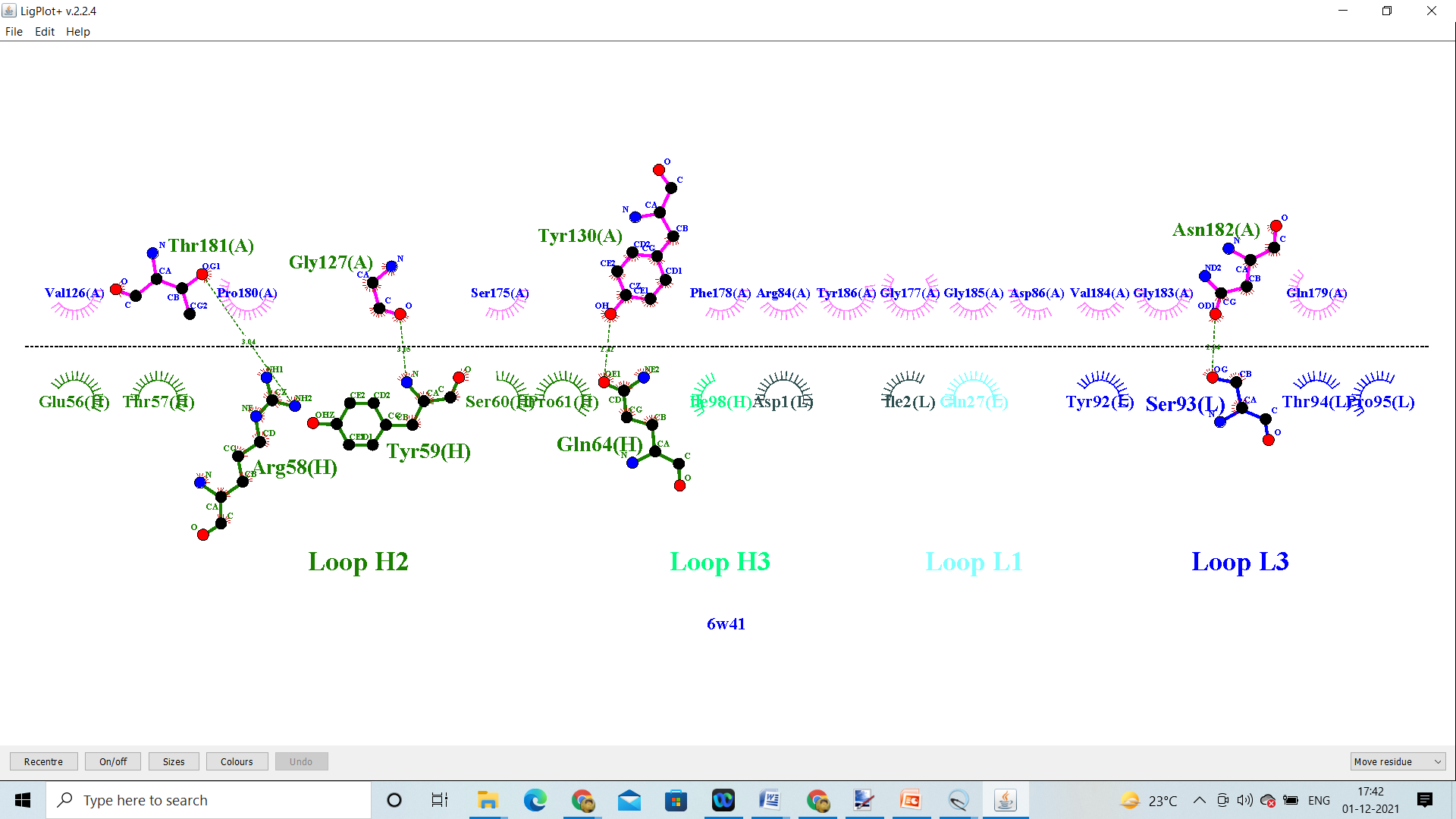

CR3022_Delta AY.3

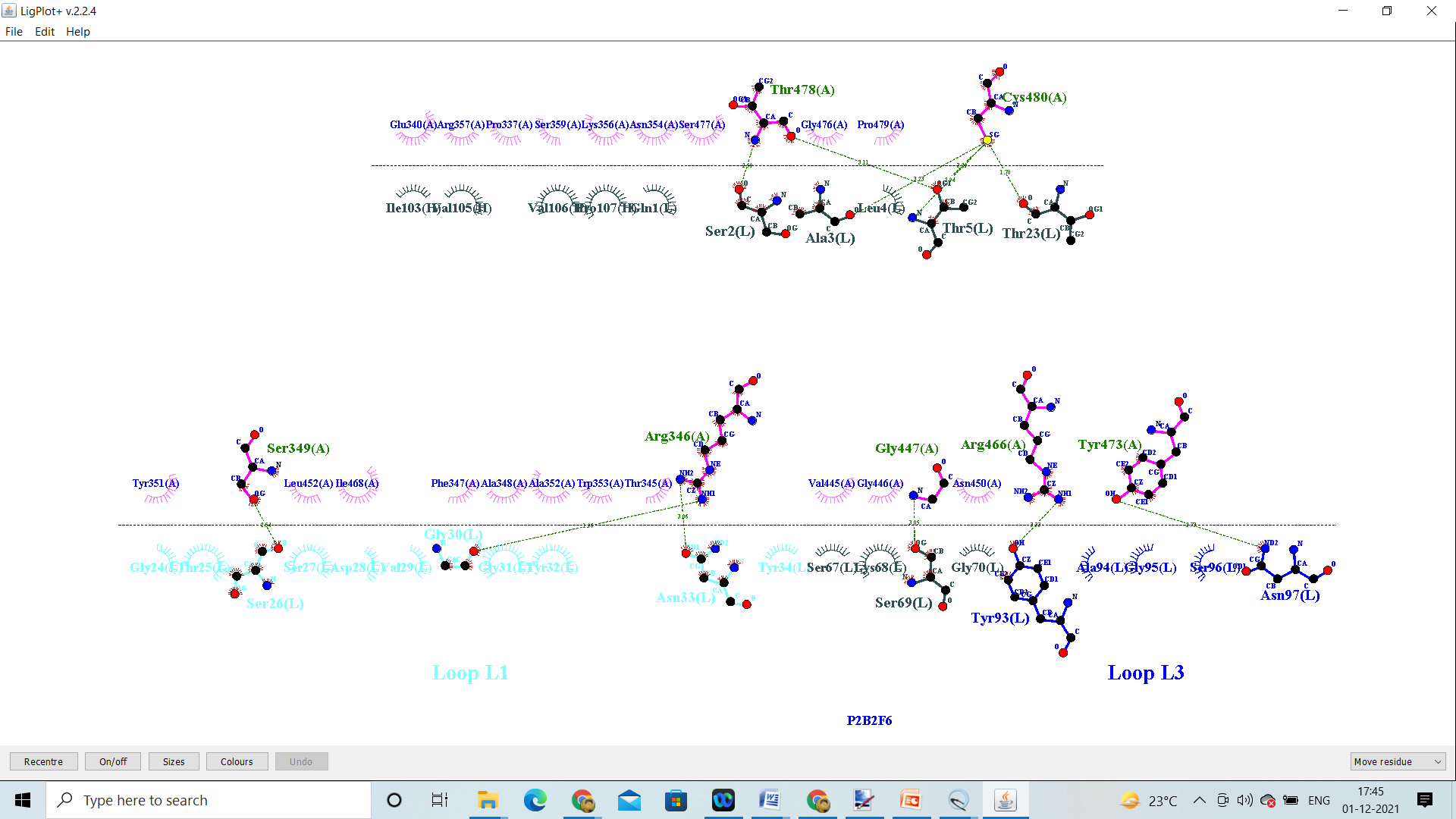

P2B2F6_Wild

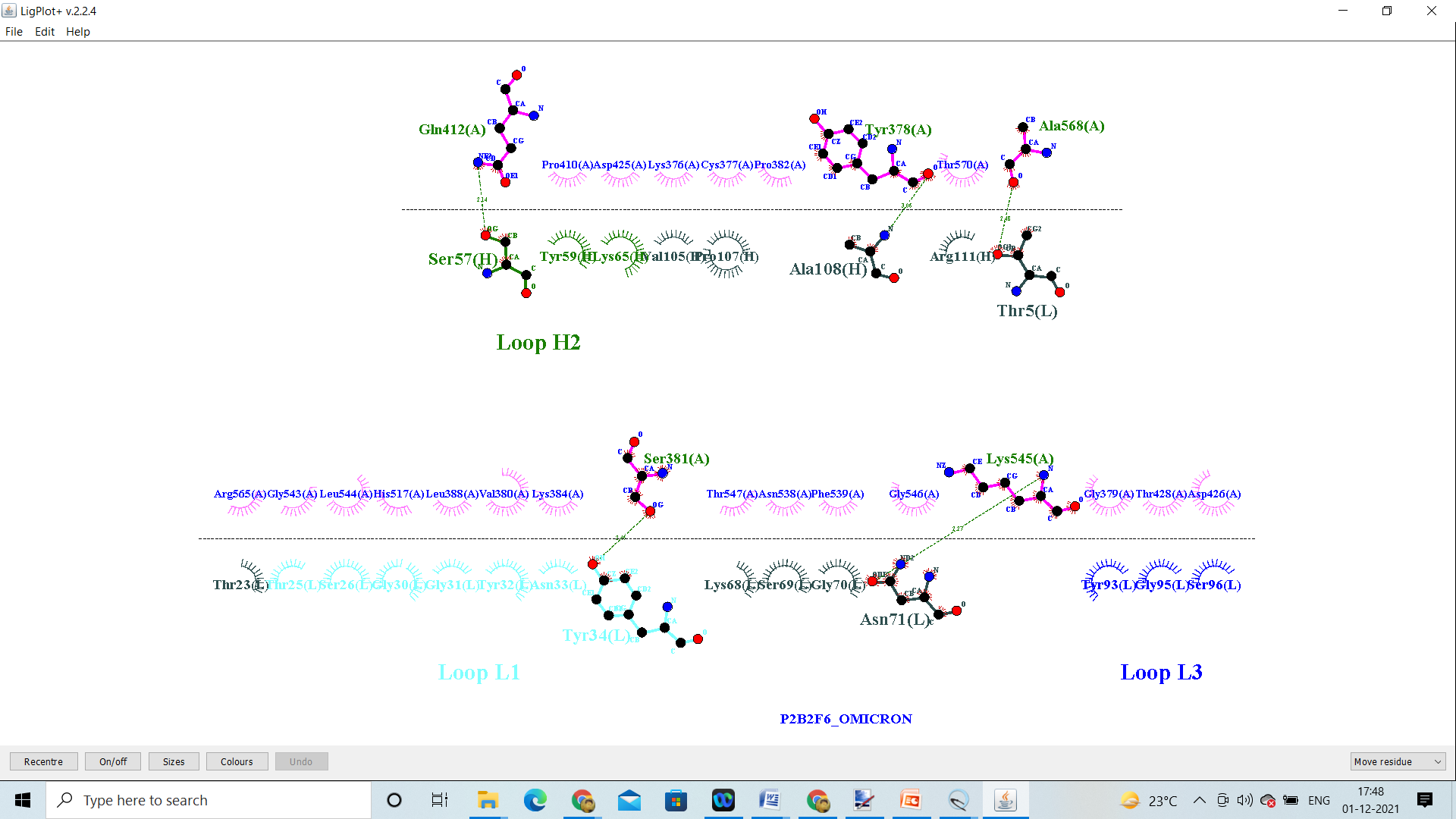

P2B2F6_Omicron

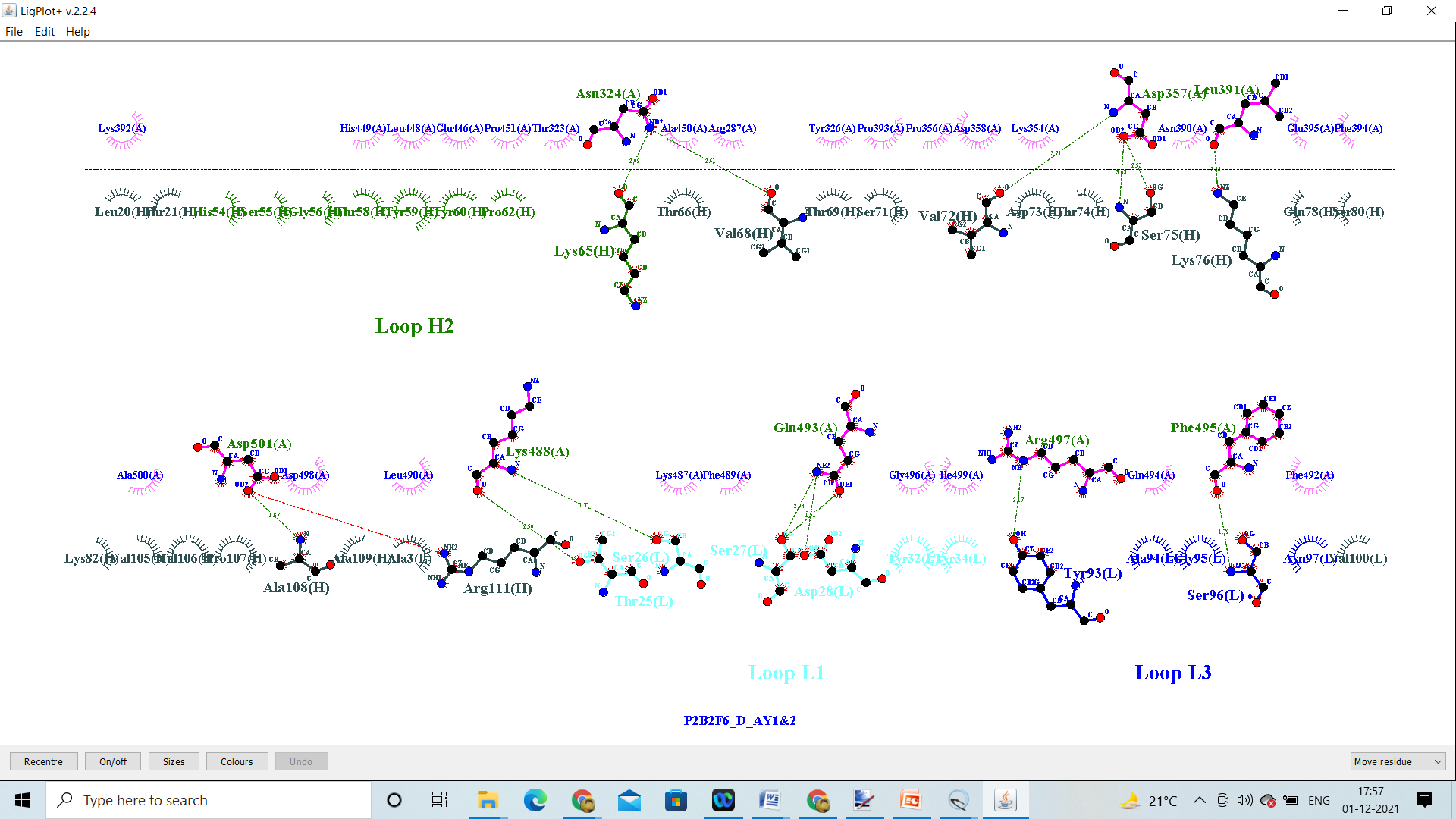

P2B2F6_Delta AY.1 &AY.2

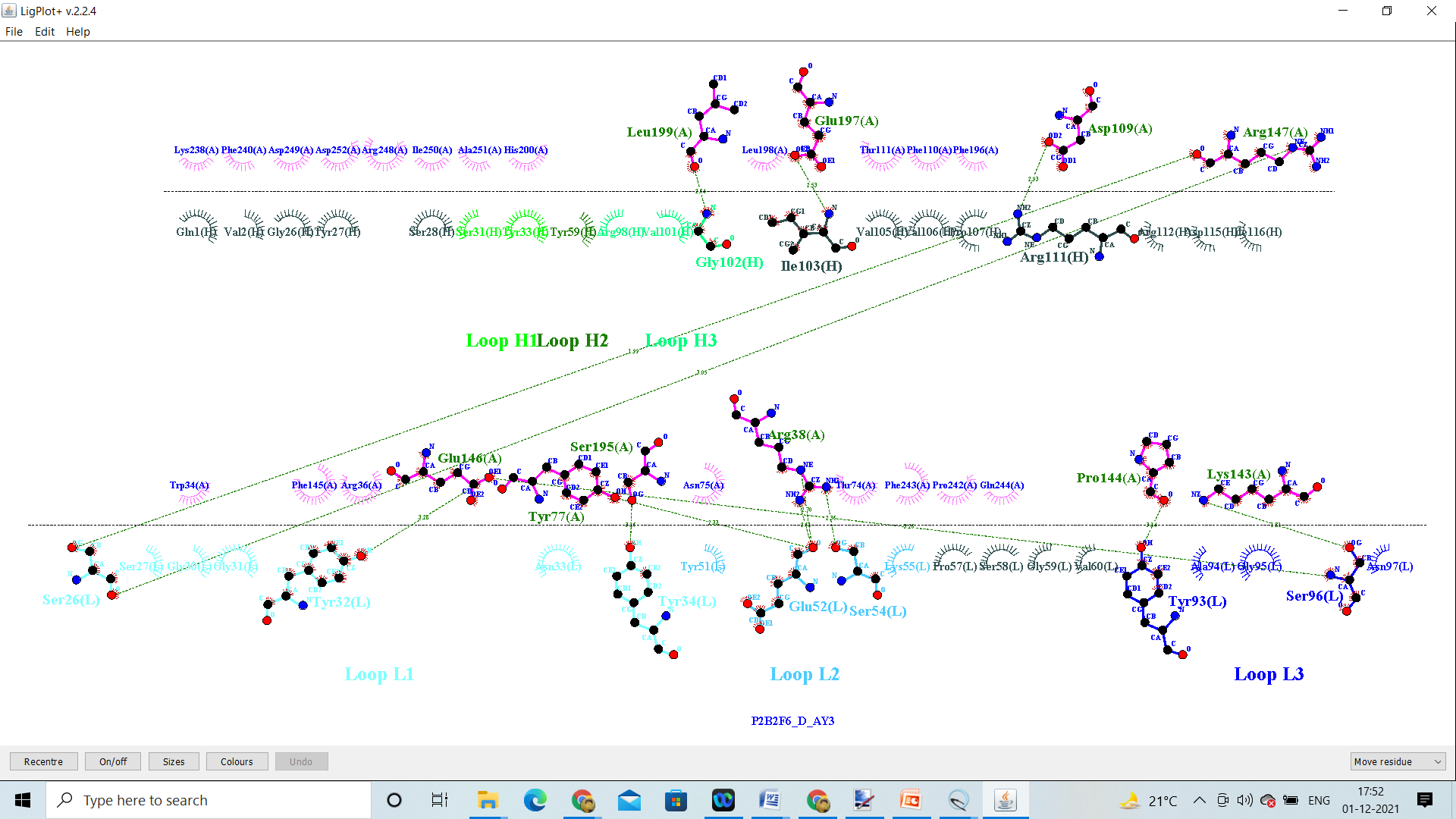

P2B2F6_Delta AY.3

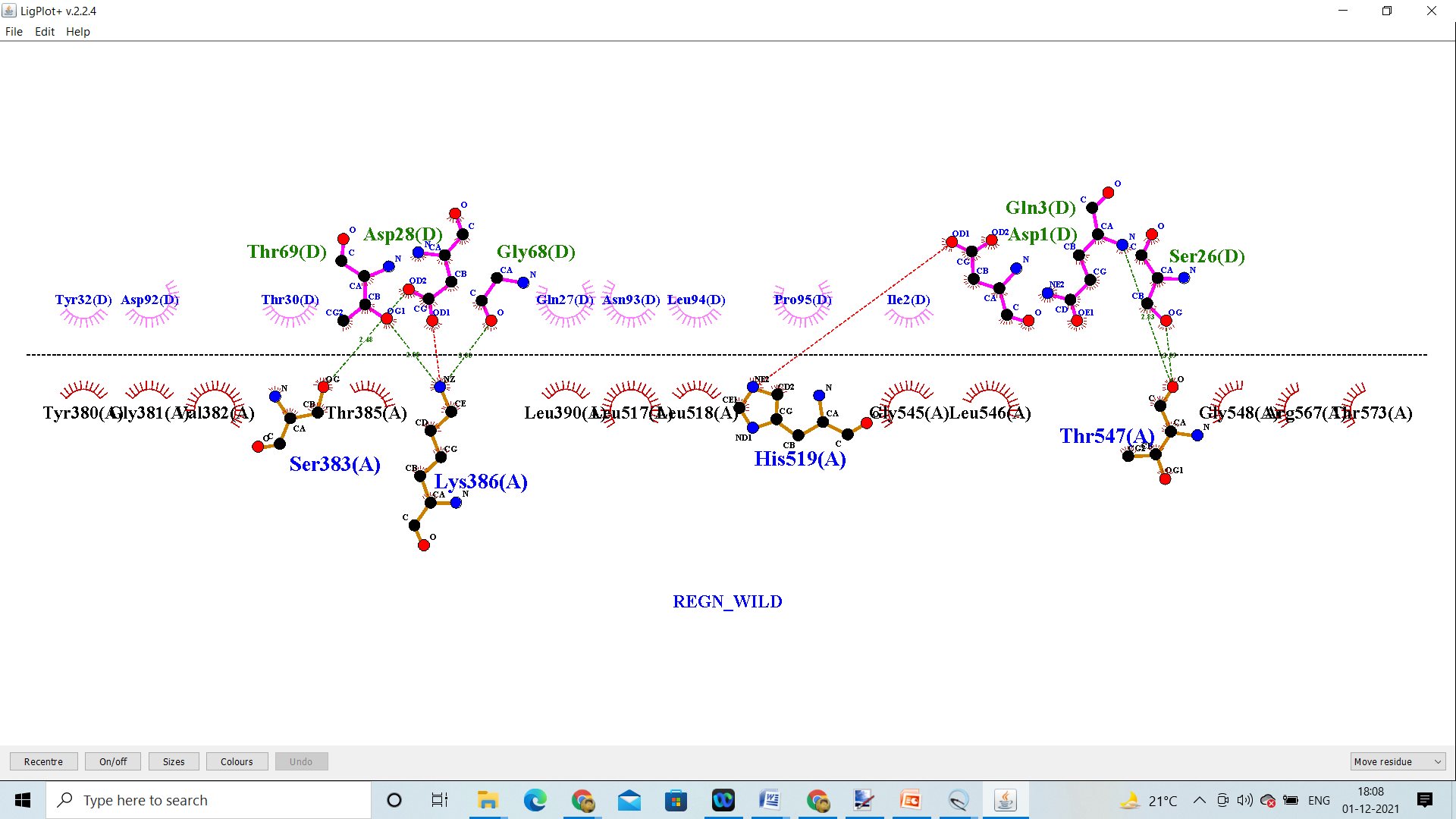

REGN_Wild

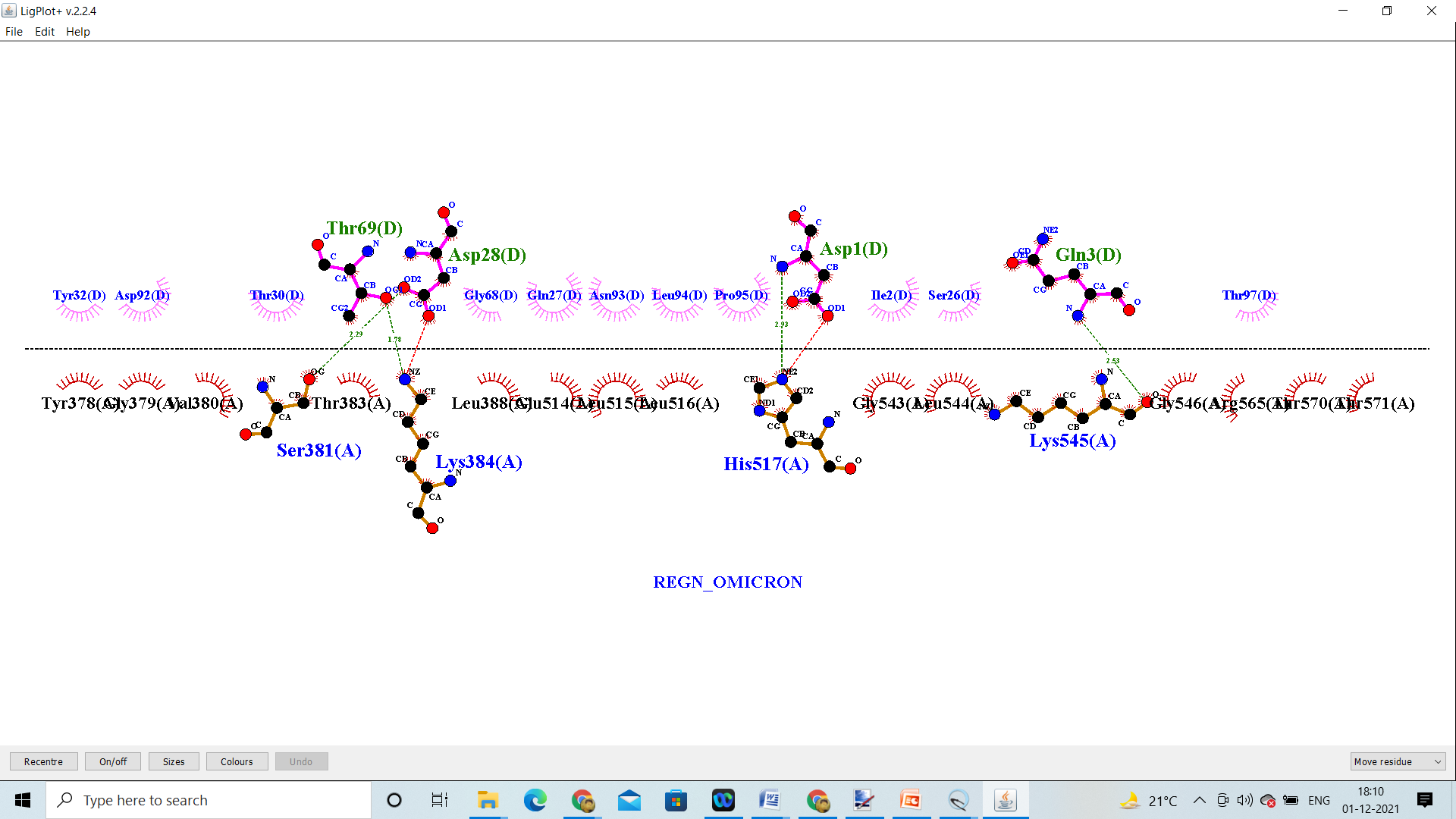

REGN_Omicron

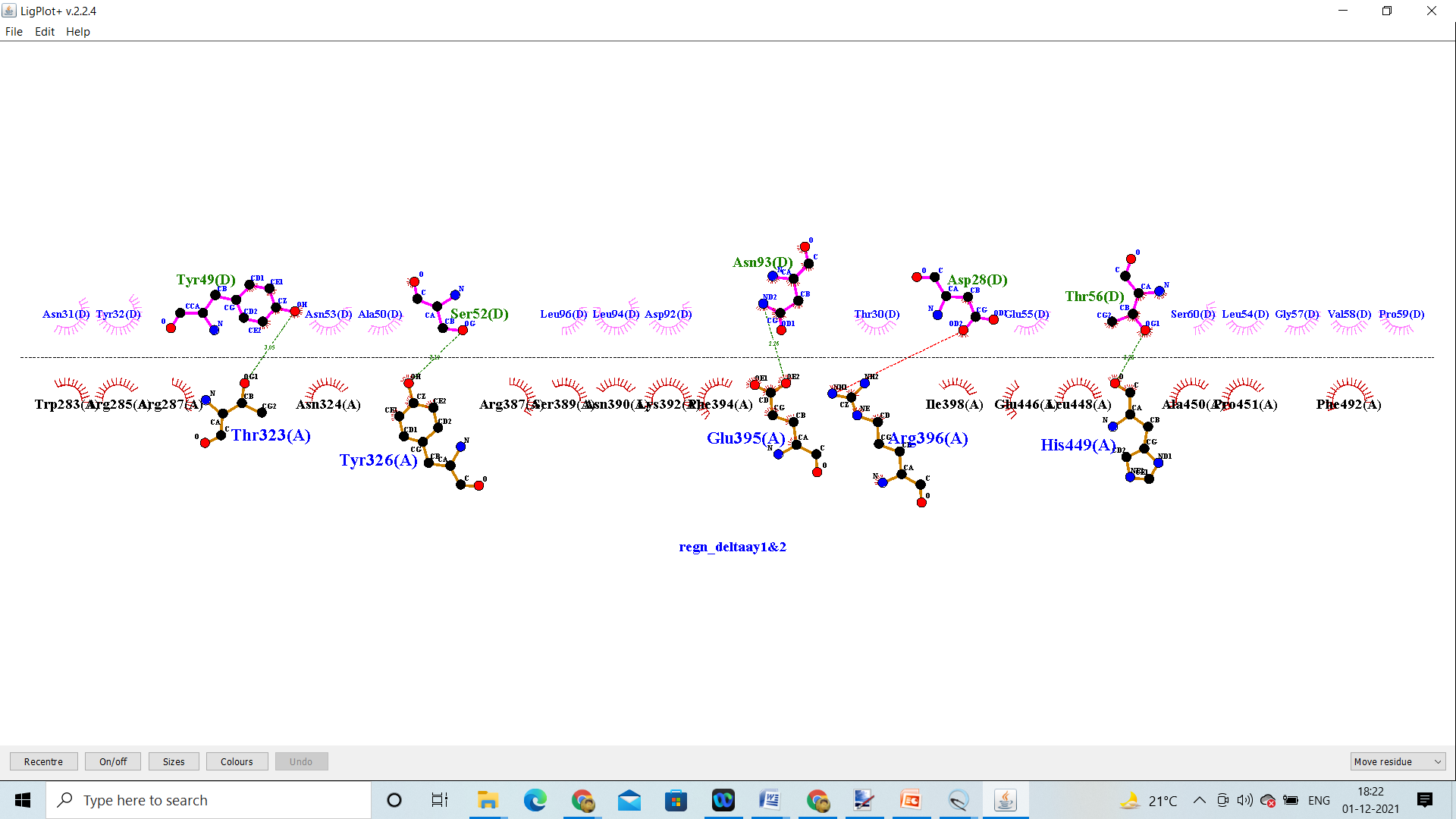

REGN_Delta AY.1 & AY.2

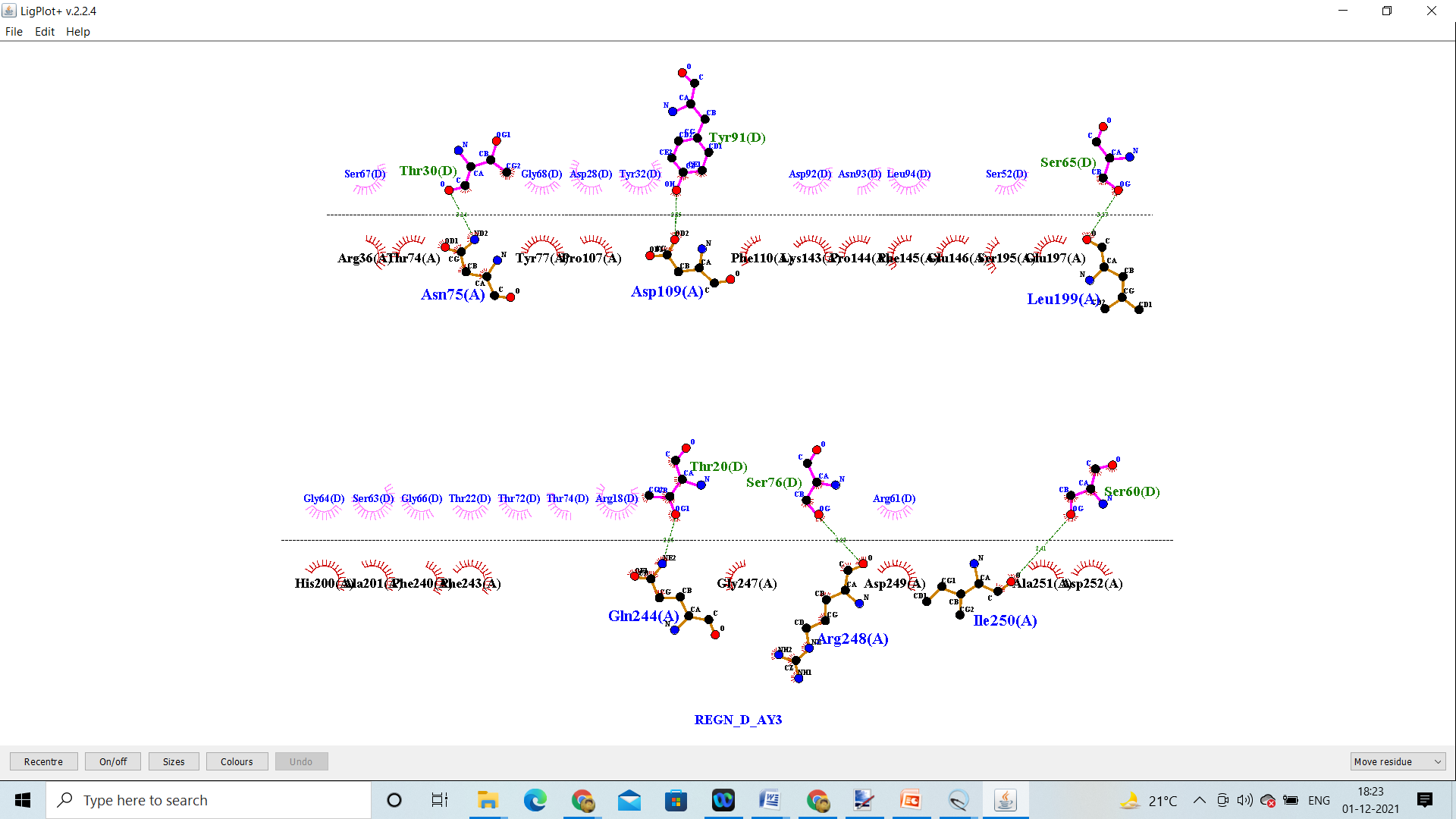

REGN_Delta AY.3

**ACE2 – RBD Variants**

Molecular interactions of ACE2 (7A97) with 3 RBD variants with structural differences in the RBD region. The green dashed lines represented hydrogen bonding, whereas the red brick represented hydrophobic interactions.

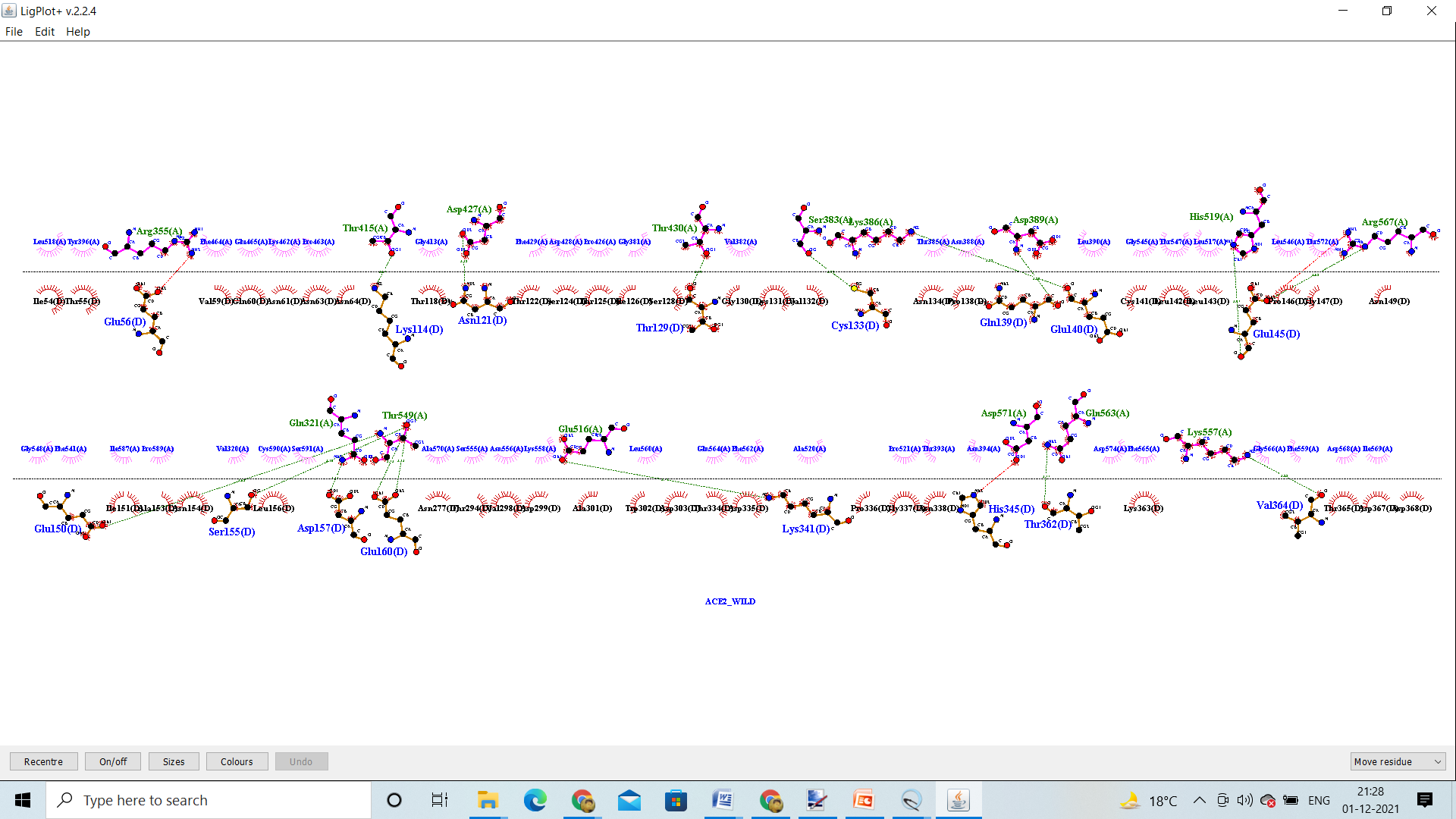

ACE2R_Wild

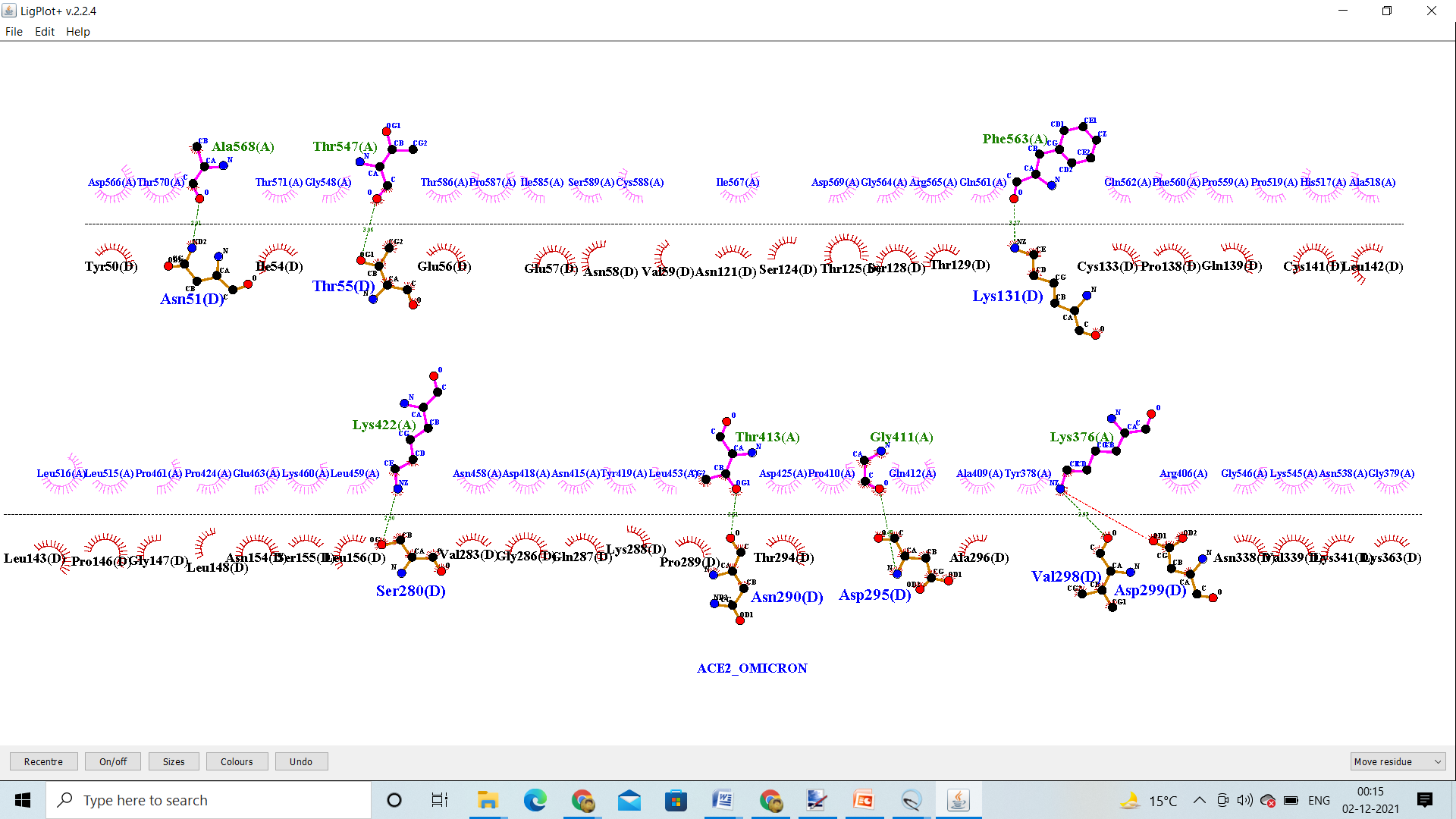

ACE2_Omicron

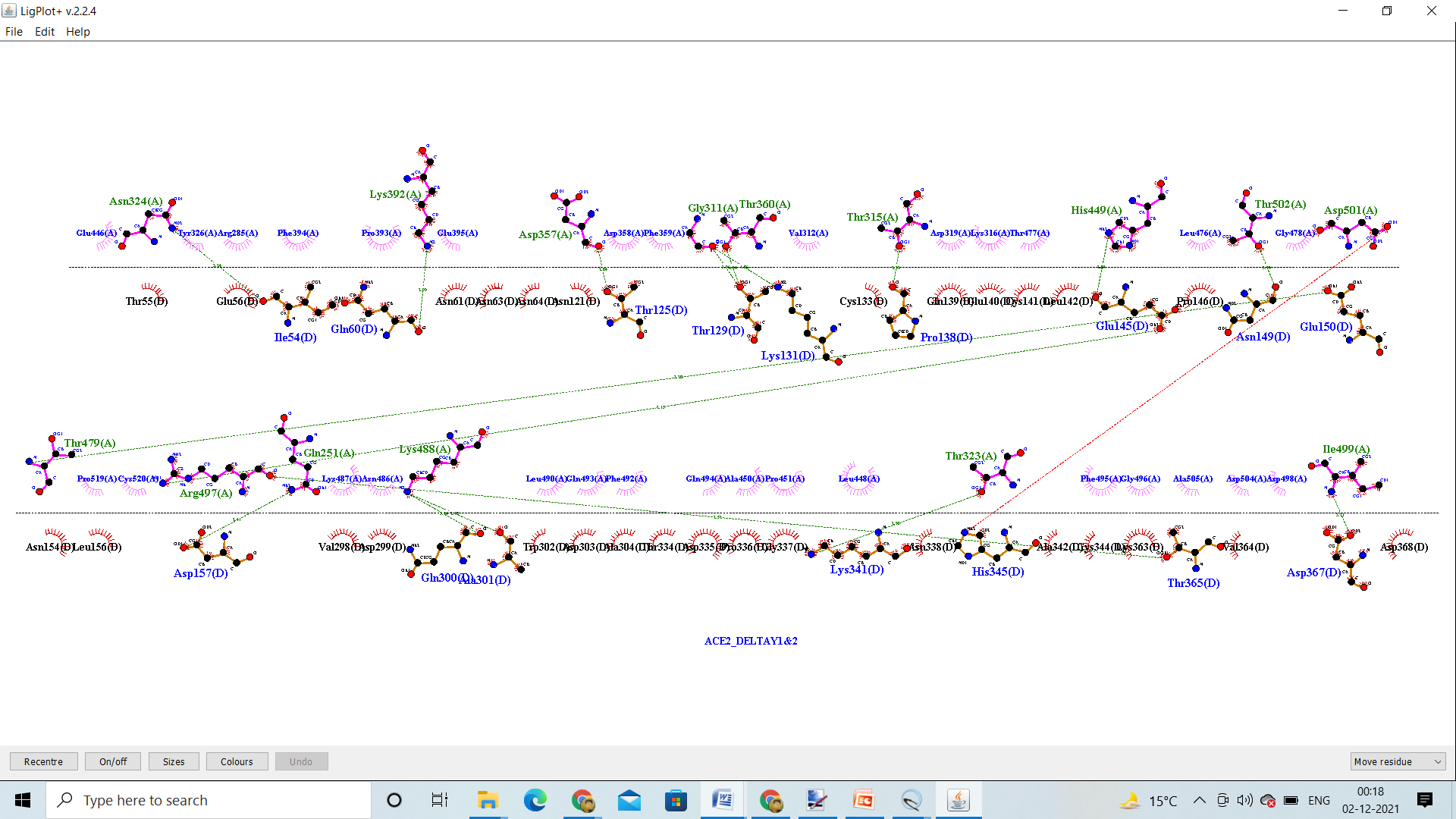

ACE2_Delta AY.1 &AY.2

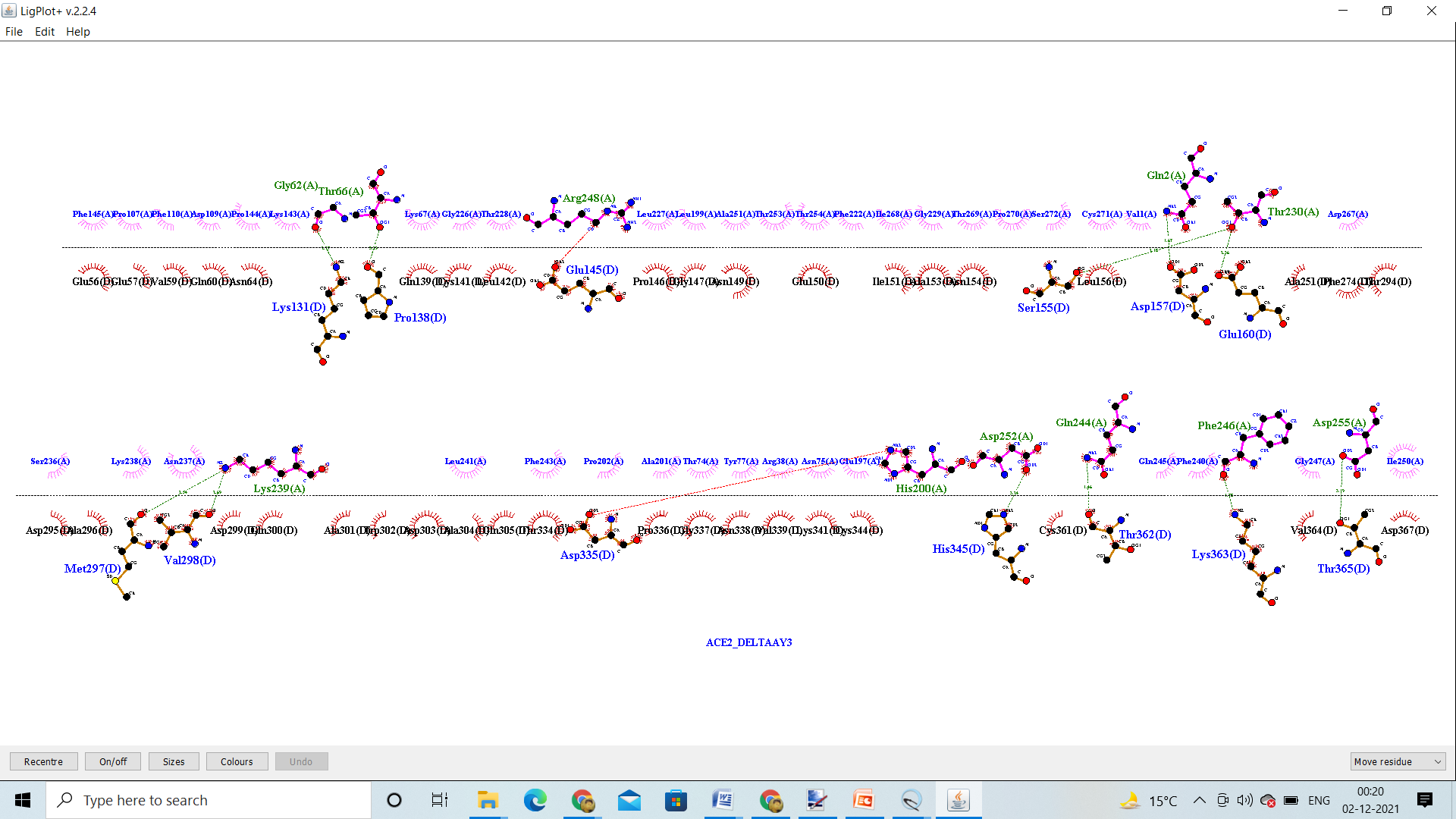

ACE2_Delta AY.3
